# Maize genetic diversity identifies moisture-dependent root-branch signaling pathways

**DOI:** 10.1101/2024.08.26.609741

**Authors:** Johannes D. Scharwies, Taylor Clarke, Zihao Zheng, Andrea Dinneny, Siri Birkeland, Margaretha A. Veltman, Craig J. Sturrock, Jason Banda, Héctor H. Torres-Martínez, Willian G. Viana, Ria Khare, Joseph Kieber, Bipin K. Pandey, Malcolm Bennett, Patrick S. Schnable, José R. Dinneny

## Abstract

Plants grow complex root systems to extract unevenly distributed resources from soils. Spatial differences in soil moisture are perceived by root tips leading to the patterning of new root branches towards available water, a process called hydropatterning. Little is known about hydropatterning behavior and its genetic basis in crops plants. Here, we develop an assay to measure hydropatterning in maize and reveal substantial differences between tropical/subtropical and temperate maize breeding germplasm that likely resulted from divergent selection. Genetic dissection of hydropatterning confirmed the regulatory role of auxin and revealed that the gaseous hormone ethylene acts to locally inhibit root branching from air-exposed tissues. These findings demonstrate the crop relevance of hydropatterning and establish its genetic basis.

## Main Text

Crop production depends heavily on water, which is a resource at risk due to the predicted effects climate change will have on the duration and severity of droughts (*1*). Plant water uptake is facilitated by an intricate network of roots. Breeding for improved root access to water is a potential tool for making crops resilient to climate change (*2*). Root networks are established by branching of the primary root axis. The development of lateral root (LR) branches is highly responsive to the spatio-temporal distribution of resources like water and nutrients in soils (*3*, *4*). Plants sense micron-scale heterogeneity in water availability at their root tips with spatial differences along the root-tip circumference determining the patterning of LRs through hydropatterning (*5*, *6*) (Fig. 1A). This response may allow plants to capture water more efficiently while minimizing the metabolic cost of root growth in dry soil (*7*). Understanding the extent of phenotypic variation for this trait within breeding populations and determining its genetic basis is required for potential applications in crop improvement. Furthermore, understanding the mechanistic basis of hydropatterning illuminates how heterogeneity in moisture is sensed by organisms to enact an adaptive response.

**Fig. 1.**
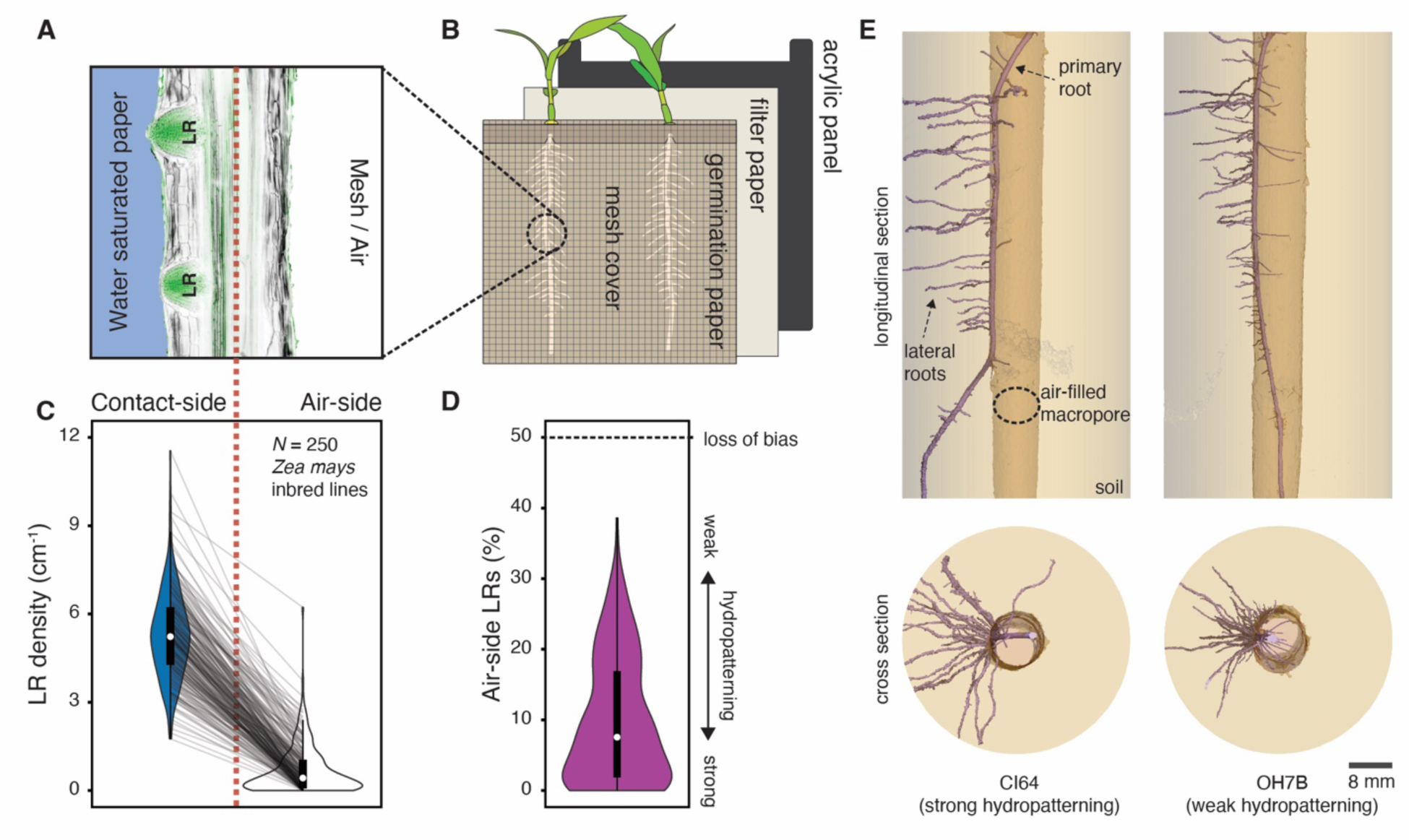
Hydropatterning responses revealed in public sector breeding lines of *Zea mays* (maize). (**A and B**) Schematic of the (A) hydropatterning response in (B) our custom-built hydropatterning assay. Primary roots of maize seedlings are grown in a vertical position along moist paper while being prevented from growing off the paper by a mesh cover. Lateral root (LR) primordia are preferentially induced towards the water-saturated paper (contact-side) and suppressed on the air-exposed side (air-side). Longitudinal cross-section of maize root (B73 inbred) stained with Calcofluor White (cell walls; gray) and SYBR Green (LRs; green). (**C and D**) Distribution of (C) contact-(blue) and air-side (white) LR densities from 250 maize inbred lines characterized using the hydropatterning assay and calculated (D) percent air-side LRs (purple). Each inbred line is represented by its median value (*n* = 1-3 seedlings/inbred) (data S2). Gray lines connect corresponding inbred lines. Population median (white circles). (**E**) X-ray Computed Tomography showing LR patterning on primary roots of strong (CI64) and weak (OH7B) hydropatterning inbred lines grown through an air-filled macropore in soil.

### Hydropatterning in domesticated maize

Here we investigated hydropatterning in the cereal crop species *Zea mays* (maize), which constitutes a major source of calories worldwide. To capture the phenotypic diversity of hydropatterning, we developed a germination paper-based assay that creates a controlled gradient of water availability across the circumference of the growing primary root (Fig. 1B and fig. S1). Simultaneous characterization of a diverse set of 250 maize inbred lines (data S1) from the Goodman-Buckler association panel allowed us to cover the majority of genetic diversity present in current public-sector breeding programs (*8*).

We observed for all tested maize inbred lines that primary roots formed LRs preferentially on the side in contact with the water-saturated germination-paper (contact-side), which is consistent with the inductive effect of water availability previously observed in the maize reference inbred B73 and in other species (*5*) (Fig. 1C, data S2). Nevertheless, a substantial proportion of all surveyed maize inbred lines developed air-side LRs as well, resulting in an observed phenotypic range of 0 - 39% air-side LRs across all 250 inbred lines (Fig. 1D). This survey revealed that a larger portion of maize inbred lines exhibit weakened hydropatterning than previously indicated (*9*), and highlighted the potential use of this trait to understand how quantitative genetic variation contributes to root architecture.

Prior research using the B73 maize inbred found that moisture availability cues, which determine the patterning of LRs, were perceived within the first 5-6 mm from the root tip (*9*). This location is well before LR founder cell divisions occur (*10*). Using additional inbred lines, we verified that pre-emerged LR primordia exhibited a similar distribution between contact- and air-sides compared to post-emergence LRs in strong and weak hydropatterning inbred lines (fig. S2, A and B). This provides further evidence that hydropatterning primarily acts at the LR founder cell patterning stage rather than at later stages.

We next examined how the observed variation in hydropatterning correlates with root architecture in soil conditions. In nature, previous root growth or burrowing invertebrate activity such as earthworms commonly create macropores, which are large air-spaces in the soil matrix. Using artificial macropores, we tested how LRs were patterned when primary roots were grown through them. Strong hydropatterning inbred lines initiated their LRs preferentially towards the side of the root in contact with soil, as observed by microscale X-ray Computed Tomography (Fig. 1E). In contrast, weak hydropatterning inbred lines displayed a reduced bias with more LRs growing into the air-filled macropore. Quantification confirmed these observations across multiple strong and weak hydropatterning inbred lines (fig. S2, C and D) and demonstrates that our hydropatterning assay generates reproducible phenotypes that are translatable to soil conditions.

To explore how variation in hydropatterning relates to other phenotypic traits of field-grown maize plants, we performed a correlation analysis with 64 compiled trait sets measured from field-grown plants (*11*). We found that both air-side LR density and percent air-side LRs correlate significantly with root crown depth and the number of nodes with brace roots (fig. S3, A to C and data S3). Weaker hydropatterning inbred lines generally exhibited more shallow root systems with fewer brace roots according to data collected by two studies from Iowa (fig. S3, D to F) (*12*, *13*). This suggests that more efficient placement of root branches towards water may improve the ability of root systems to attain greater depths, possibly due to the metabolic savings achieved by limiting branching in dry soil (*14*). Importantly, no significant correlations were observed with contact-side LR density, suggesting that this trait has less relevance to the in-field root architecture traits measured.

### Phenotypic diversity and genetics of hydropatterning

It has been shown that domestication and local environmental conditions established selective pressures on maize landraces that have determined the current structure of phenotypic and genotypic diversity in modern inbreds (*15–17*). Therefore, we explored whether hydropatterning has been subjected to selection and analyzed the associated current genetic variation in maize.

### Variation and selection across maize breeding subpopulations

We reassessed population structure and assigned subpopulations for all phenotyped inbred lines that had matching whole-genome sequencing data available (*n* = 231). In accordance with previous research, we assigned inbred lines to a mixed group when subpopulation identity was less than 80% (*8*) (Fig. 2A and data S4). While variation in contact-side LR density was uniform across all subpopulations, inbred lines with higher air-side LR density and percent air-side LRs were predominantly associated with the non-stiff stalk/mixed groups (Fig. 2, B to D). This led to significantly weaker hydropatterning (more air-side LRs) for the large temperate non-stiff stalk group compared to the large tropical/subtropical group or the smaller temperate stiff stalk group (Fig. 2E and fig. S4A).

**Fig. 2.**
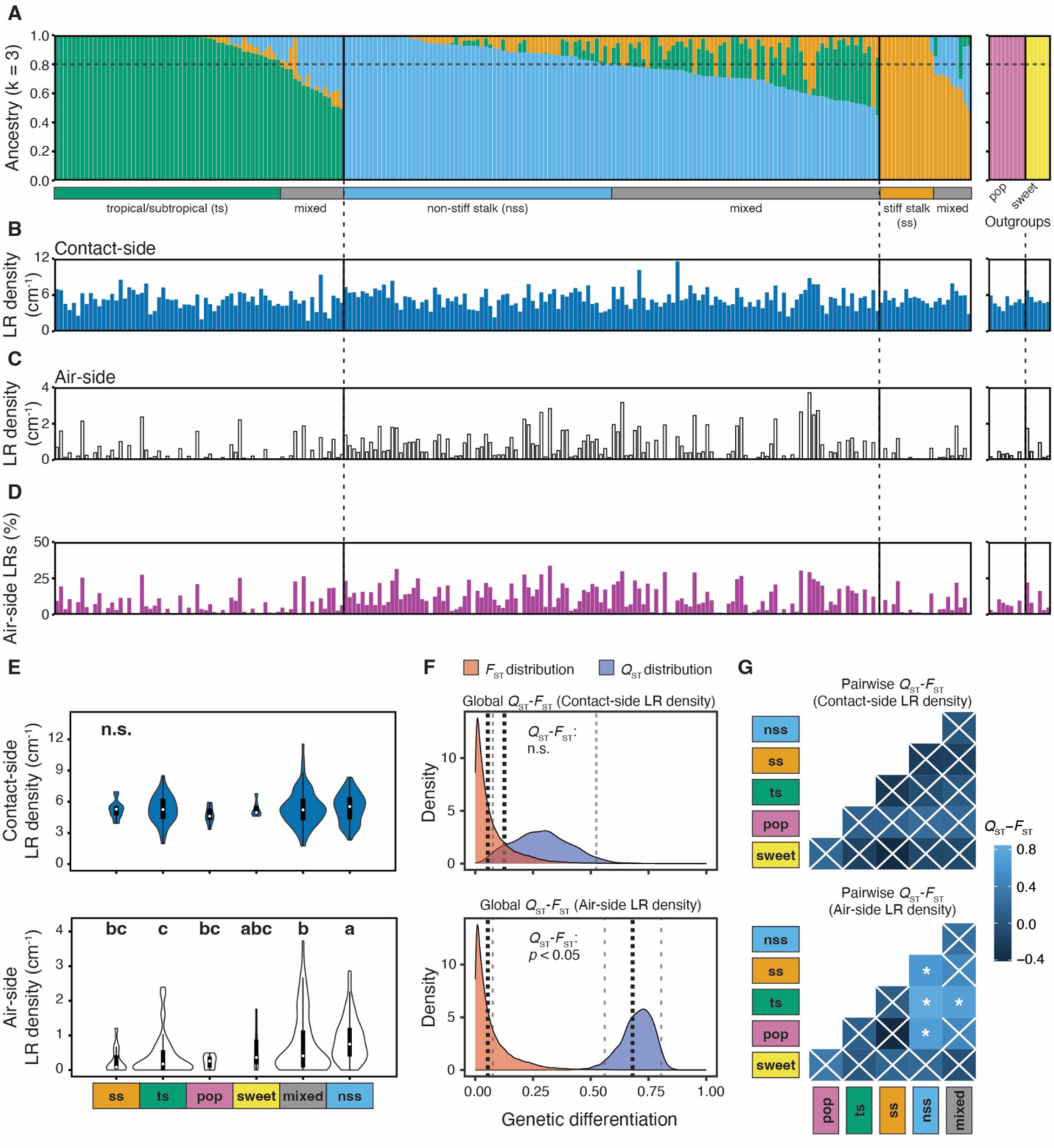
Differences in hydropatterning across breeding subpopulations may have been caused by divergent selection. **(A - D)** Population structure and hydropatterning traits of 231 maize inbred lines (19 inbred lines from original population of *N* = 250 were excluded due to missing genotypic data). (A) Ancestry components and subpopulation assignments. Inbred lines < 80% group identity = “mixed”. Popcorn (pop) and sweet corn (sweet) groups were defined a priori. Phenotypic data of (B) contact-side and (C) air-side LR density, and (D) percent air-side LRs are shown as the median value for each inbred line (*n* = 1-3 seedlings/inbred) (data S2). (**E**) Subpopulation comparisons of contact-side (top) and air-side (bottom) LR density. Violin plot area adjusted for number of inbred lines / subpopulation. Letters denote significant differences between subpopulations (*p* ≤ 0.05, Kruskal–Wallis and Dunn’s post hoc tests); n.s., no significant differences. (**F**) Population-wide comparison of *F*_ST_ (fixation index) and *Q*_ST_ (genetic differentiation in regards to a quantitative trait) distributions for contact-side (top) and air-side (bottom) LR density. Black dotted lines (means), gray dashed lines (95% confidence intervals). (**G**) Subpopulation pairwise *Q*_ST_ - *F*_ST_ comparisons. Asterisks denote significant differences between *Q*_ST_ and *F*_ST_ (Benjamini & Hochberg-adjusted: * *p* ≤ 0.05, ** *p* ≤ 0.01, *** *p* ≤ 0.001); white crosses, not significant. Number of inbred lines in each subpopulation across all panels: *n*_ts_ = 53, *n*_nss_ = 63, *n*_ss_ = 14, *n*_mixed_ = 88, *n*_pop_ = 9, *n*_sweet_ = 6.

To test whether this divergence in hydropatterning was best explained by neutral evolution or selection, we compared quantitative genetic differentiation with regards to contact- or air-side LR density (*Q*_ST_) to the population genetic differentiation due to genetic structure (*F*_ST_). We found that *Q*_ST_ and *F*_ST_ distributions overlapped for contact-side LR density, which suggests that genetic differentiation for this trait occurred through neutral evolution. Conversely, *Q*_ST_ was significantly in excess of *F*_ST_ for air-side LR density, as well as percent air-side LRs, indicating divergence through differential selection for these hydropatterning traits (Fig. 2F, fig. S4B, and data S5). Pairwise comparisons between subpopulations show the largest *Q*_ST_ - *F*_ST_ differences between the non-stiff stalk and tropical/subtropical groups, suggesting that divergence in selective pressures for hydropatterning occurred predominantly after the split between tropical/subtropical and temperate germplasm (Fig. 2G, fig. S4C, and data S5). Stronger hydropatterning in the tropical/subtropical subpopulation may have resulted from selection for drought tolerance, among other abiotic stress factors, during breeding (*18*, *19*), while weakened hydropatterning in the temperate non-stiff stalk subpopulation could be the result of relaxation in selection pressures on efficient water uptake in temperate environments. Although pronounced *Q*_ST_ > *F*_ST_ differences for air-side LR density and contrasting observations for contact-side LR density support our hypothesis, it should be noted that inbreeding may inflate *Q*_ST_ estimates (*20*).

### Genetic architecture of hydropatterning in domesticated maize

Previous work in *Arabidopsis thaliana* (Arabidopsis) has shown that the auxin hormone pathway promotes branching on the contact side of roots during hydropatterning. Mutants that disrupt auxin biosynthesis and polar transport are known to weaken hydropatterning (*5*). Furthermore, it was found that the auxin-response transcription factor AUXIN RESPONSE FACTOR 7 (ARF7) is sumoylated in cells on the air-side of roots, promoting binding to the repressor protein INDOLE-3-ACETIC ACID INDUCIBLE 3 (IAA3) which blocks initiation of LR founder cells (*21*). To identify novel loci and associated genes with relevance to maize, we conducted Genome and Transcriptome Wide Association Studies (GWAS/TWAS).

TWAS (*13*) identified nine genes whose expression in germinating seedling roots (*22*) is significantly associated with differences in air-side LR density (Fig. 3A and data S6). Among these TWAS-genes, we found *Zm00001eb211770*, a maize ortholog of *AUXIN RESISTANT 1* (*ZmAXR1*). In Arabidopsis, it has been shown that AXR1 promotes auxin responses through its function as a subunit of RUB1, an activating enzyme of the SCF-complex that is associated with protein-degradation and auxin perception (*23*). Correlation analysis suggests a positive relationship between *ZmAXR1* gene expression and air-side LR density (fig. S5A). To analyze the origin of the variation in *ZmAXR1* gene expression, we mapped the associated expression Quantitative Trait Loci (eQTL). This analysis revealed several highly significant expression-associated Single Nucleotide Polymorphisms (e-SNP) for *ZmAXR1* (fig. S5B, data S7 and 8). The most significant e-SNP, Chr5_2705946, is co-localized with *ZmAXR1* itself, indicating that cis-acting regulatory variation may explain the variation in *ZmAXR1* gene expression associated with air-side LR density. Input data for the TWAS suggested expression of *ZmAXR1* in root tips of most inbred lines (fig. S6A). This was confirmed through *in situ* detection of *ZmAXR1* transcripts in root tips of maize inbred line B73 by Hybridization Chain Reaction (HCR) (fig. S6, B and C). These results indicate that auxin regulation and related processes play a significant role in hydropatterning for maize roots, as previously demonstrated in Arabidopsis.

**Fig. 3.**
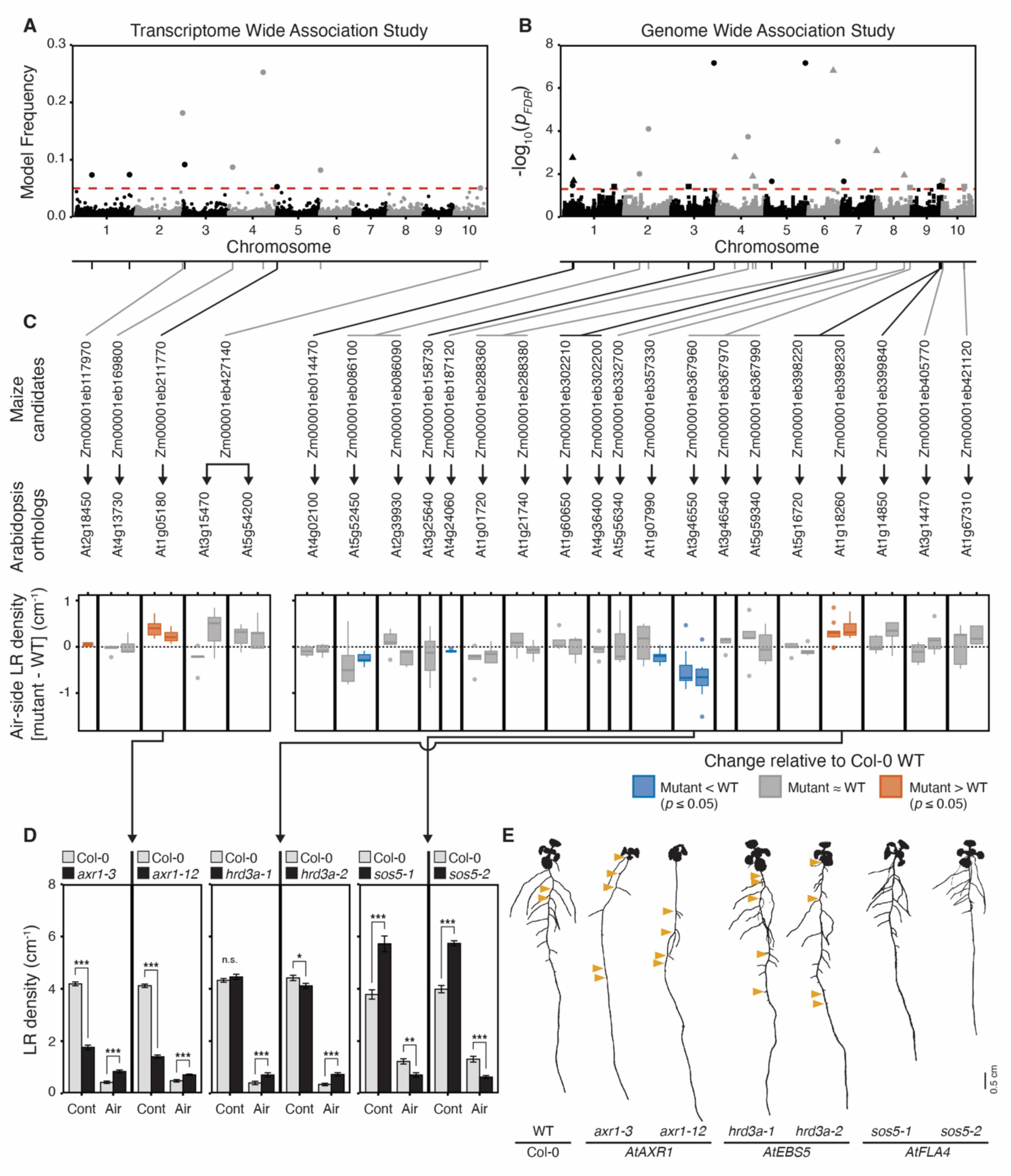
Genetic control for hydropatterning revealed in maize and validated in Arabidopsis. **(A and B)** Manhattan plots of Transcriptome Wide Association Study (TWAS) and Genome Wide Association Studies (GWAS) for air-side lateral root (LR) density in maize. (A) TWAS used gene expression data from maize root tips (*22*). Significance threshold: model frequency = 0.05 (red dashed line). (B) GWAS shows smallest *p*-value for each Single Nucleotide Polymorphism (SNP) across three Minor Allele Frequency cutoffs ≥ 0.4% (solid circles), ≥ 2.2% (solid triangles), and ≥ 4.4% (solid squares). FDR-adjusted significance threshold: *p* = 0.05 (red dashed line). Gray and black lines connect SNPs and associated candidate genes. (**C**) Validation of maize candidate genes using mutants of corresponding gene orthologs in Arabidopsis. Air-side LR density of mutants are shown relative to Col-0 wild-type (WT). Fill color denotes significant differences (Paired Student’s t-test, *p* ≤ 0.05) mutants > WT (orange), mutants < WT (blue), not significant (gray) *n* = 5 - 10 plates per mutant (5 WT & 5 mutant seedlings/plate). (**D**) Comparisons of contact-side (Cont) and air-side (Air) LR densities between Col-0 wild-type (gray) and mutants for *AtAXR1*, *AtEBS5*, and *AtFLA4* (all mutants black). *n* = 10 plates per mutant (5 WT & 5 mutant seedlings/plate). Asterisks denote significant differences (Paired Student’s t-test: * *p* ≤ 0.05, ** *p* ≤ 0.01, *** *p* ≤ 0.001). Bar height (mean), error bar (standard error). (**E**) Binary images of 11 day-old seedlings. Orange triangles mark air-side LRs. Scale bar = 0.5 cm. Dilation was used on images to improve visibility.

In parallel, GWAS (*24*, *25*) identified 30 unique Trait-Associated SNPs (TASs) for air-side LR density (Fig. 3B and data S9), suggesting that variation in hydropatterning is controlled by numerous loci in maize. In almost all cases, higher air-side LR density was associated with the minor allele of TAS, except for TAS Chr8_143668219. This supports our hypothesis that weakening of hydropatterning may have been caused by relaxation of selection, since selection typically constrains the occurrence of genetic variants (*26*). We identified 40 genes within 20-kb windows centered on the TAS. In cases where no gene was located within these windows, the next closest gene was included (data S10). These genes are candidates that may control air-side LR density in maize.

### Validation of maize candidate genes through genetic analysis of orthologues in Arabidopsis

To explore the molecular basis of variation in gene function associated with the maize candidate genes from TWAS and GWAS, we identified their gene orthologs in Arabidopsis and screened available mutant lines for hydropatterning defects using a gel-based assay (fig. S7A and data S11 and 12). We identified seven genes for which at least one mutant allele showed a significant change in air-side LR density and percent air-side LRs (Fig. 3C and fig. S7B). For three of these genes, two independent mutant alleles both showed significant, matching changes in air-side LR density, confirming their association with hydropatterning (Fig. 3, D and E)

Mutants *axr1-3* and *axr1-12* of *AtAXR1* showed significant increases in air-side LR density and percent air-side LRs compared to the wild-type (WT), while contact-side LR density dropped significantly (Fig. 3D). This defect is similar to the phenotype of other auxin-pathway mutants in Arabidopsis (*5*, *21*) and confirms that the auxin hormone pathway plays an important role in promoting the bias in LR development towards the moisture-contacting side of the root both in Arabidopsis and maize.

Similarly, mutants *hrd3a-1* and *hrd3a-2* of *EMS-MUTAGENIZED BRI1 SUPPRESSOR 5* (*AtEBS5*), an ortholog of GWAS candidate *Zm00001eb398230* (*ZmHRD3A*), showed significant increases in air-side LR density and percent air-side LRs compared to the WT (Fig. 3D). However, in contrast to mutants of *AtAXR1*, *hrd3a-1* and *hrd3a-2* showed no, or a relatively small, change in contact-side LR densities, respectively. These data suggest that *HRD3A* may primarily function in the suppression of air-side LR development. In maize, *ZmHRD3A* was discovered as one of three candidate genes associated with TAS Chr9_147576641 (data S10). Mutants of *At5g16720*, an ortholog of *Zm00001eb398220* associated with the same TAS, did not affect hydropatterning (fig. S7B). In Arabidopsis, *AtEBS2* encodes a subunit of the Hrd1 complex and functions in the endoplasmic reticulum-associated protein degradation (ERAD) pathway (*27*). This pathway targets misfolded receptor-like kinases and glycosylated proteins (*28*), suggesting that hydropatterning may rely upon proteins acting at the plasma membrane that are also targets of ERAD. Both analysis of gene expression data from root tips, as well as *in situ* detection by HCR showed that *ZmHRD3A* is expressed at the root tip and, hence, is able to participate in the hydropatterning response (fig. S6, A and D).

In contrast to *axr1* and *hrd3a,* mutants *sos5-1* and *sos5-2* of *FASCICLIN-LIKE ARABINOGALACTAN-PROTEIN 4* (*AtFLA4*), an ortholog of GWAS candidate *Zm00001eb367960* (*ZmFLA4*), showed significant strengthening of hydropatterning with decreases in air-side LR density and percent air-side LRs compared to WT (Fig. 3D). This change was accompanied by a significant increase in contact-side LR density and a significant reduction in primary root length (Fig. 3E and fig. S8A). Taking this reduction of primary root length into account, *sos5-1* and *sos5-2* both showed a highly significant decrease in the total number of emerged air-side LRs per seedling but only a small or no increase in total contact-side LRs per seedling (fig. S8B). Thus, these data indicate that *sos5* primarily suppresses air-side LRs, while changes in contact-side LR density may result from pleiotropic effects on root length. In Arabidopsis, the mutant *sos5-1* was identified for its defects in growth on saline media (*29*).

While the primary root phenotype under salinity and ionic stress has been studied extensively (*30*), no reports have described a LR phenotype. *AtFLA4* belongs to a group of 21 fasciclin-like arabinogalactan proteins and carries two fasciclin 1 domains that allow it to interact with the extracellular cell wall matrix (*31*, *32*), which may allow it to sense extracellular cues originating from the environment. In maize, *ZmFLA4* was discovered as one of four candidate genes associated with TAS Chr8_175425640 (data S10). Orthologous Arabidopsis mutants of two other candidate genes associated with the same TAS, *Zm00001eb367970* and *Zm00001eb367990*, showed no defect in hydropatterning (fig. S7B). This provides evidence that variation associated with *ZmFLA4* may be the primary determinant of the observed variation in hydropatterning at TAS Chr8_175425640. In maize, analysis of gene expression data from roots tips, as well as *in situ* detection by HCR showed that *ZmFLA4* is expressed at the root tip and, hence, is able to participate in the hydropatterning response (fig. S6, A and E).

### Ethylene as an air-side signal mediating hydropatterning

Ethylene is a gaseous plant hormone that regulates development in response to several abiotic stresses (*33*). Accumulation of root-produced ethylene in compacted soils has been shown to serve as a signal leading to root growth inhibition (*34*). Previous research in Arabidopsis has suggested that *AtFLA4* may act in a genetic pathway regulating the synthesis of ethylene (Fig. 4A) (*30*). In this pathway, AtFLA4 acts upstream of two leucine-rich repeat receptor-like kinases, AtFEI1 and AtFEI2 (*35*). These kinases are known to interact with 1-AMINOCYCLOPROPANE-1-CARBOXYLATE SYNTHASE (ACS) 5 and 9, which are involved in the synthesis of the ethylene precursor 1-aminocyclopropane-1-carboxylate (ACC). Associated with this pathway, ETHYLENE OVERPRODUCER 1 (AtETO1) functions as a negative regulator of type-2 ACS enzymes, including ACS5 and ACS9 (*36*).

**Fig. 4.**
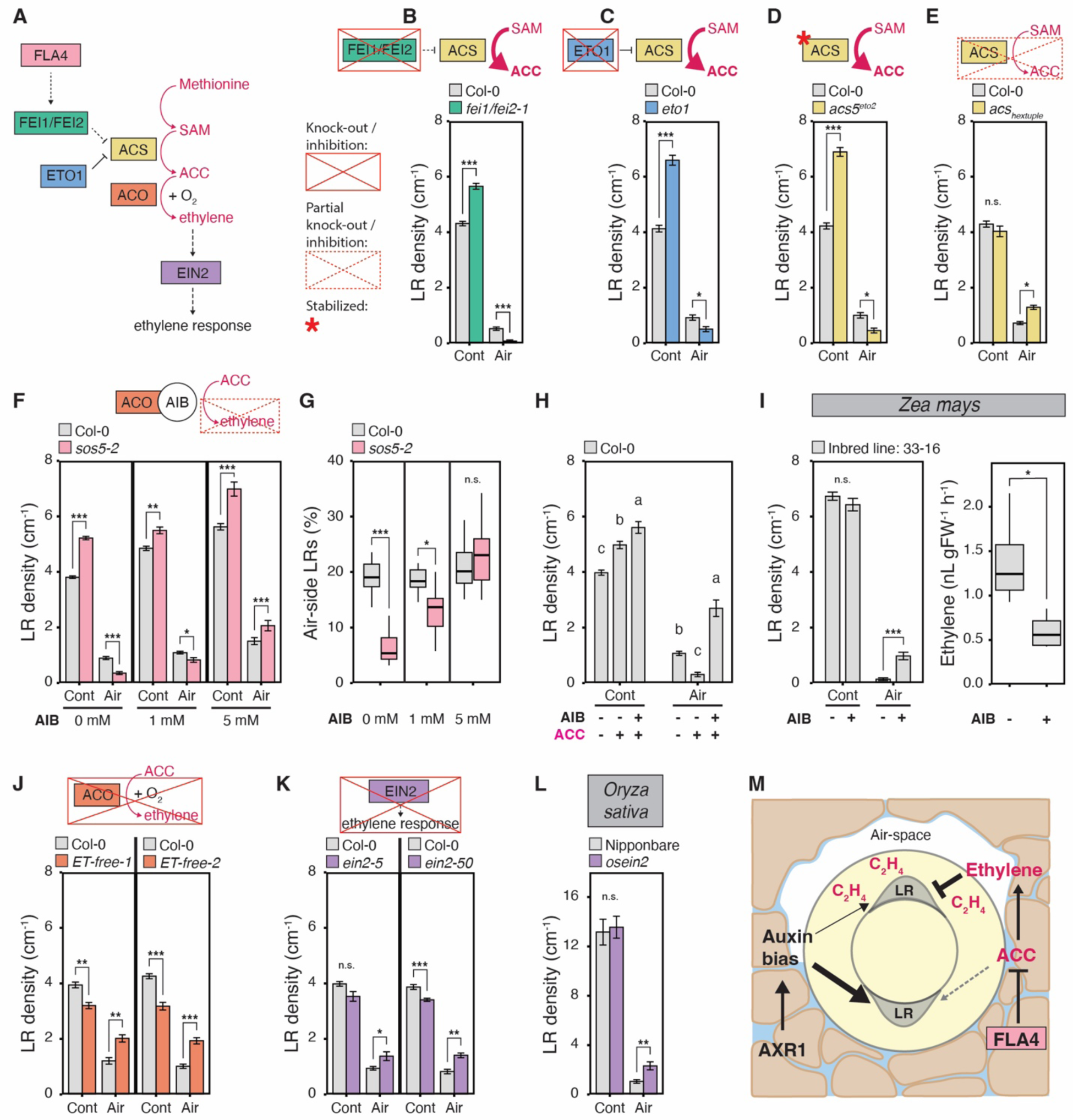
Ethylene inhibits formation of air-side lateral roots in Arabidopsis, maize, and rice. (**A**) FLA4 and the ethylene pathway in Arabidopsis (*30*). FLA4 may repress the conversion of S-Adenosyl methionine (SAM) into the ethylene precursor 1-Aminocyclopropane-1-carboxylic acid (ACC) by inhibiting ACC synthases (ACSs) via receptor kinases FEI1/FEI2. (**B - E, J, and K**) Comparisons of contact-side (Cont) and air-side (Air) lateral root (LR) densities between Col-0 (gray) and mutants *fei1/fei2-1* (green, *n* = 6 plates), *eto1* (blue, *n* = 5 plates), *acs5^eto2^* (yellow, *n* = 5 plates), *acs2-1/4-1/5-2/6-1/7-1/9-1* hextuple mutant (yellow, *n* = 5 plates), *ET-free-1* & *ET-free-2* (orange, *n* = 8-9 plates), and *ein2-5* & *ein2-50* (purple, *n* = 10 plates) in Arabidopsis. (**F - H**) Treatment of Col-0 (gray) and *sos5-2* (pink) with 2-Aminoisobutyric acid (AIB) and ACC. In (H) treatments are (-) mock control and (+) 0.05 mM ACC / 5 mM AIB. *n* = 10 plates/treatment. For Arabidopsis, asterisks denote significant differences between Col-0 and mutant (Paired Student’s t-test: * *p* ≤ 0.05, ** *p* ≤ 0.01, *** *p* ≤ 0.001), while letters denote significant differences between treatments (Student’s t-test, FDR-adjusted: *p* ≤ 0.05). (**H**) Maize inbred line “33-16” treated with (-) mock control or (+) 10 mM AIB. Asterisks denote significant differences between treatments (Student’s t-test: * *p* ≤ 0.05, ** *p* ≤ 0.01, *** *p* ≤ 0.001). (**L**) Cont and Air LR densities in wild-type rice cultivar Nipponbare (gray, *n* = 16 seedlings) and mutant *osein2* (purple, *n* = 11 seedlings). Asterisks denote significant differences between WT and mutant line (Student’s t-test: * *p* ≤ 0.05, ** *p* ≤ 0.01, *** *p* ≤ 0.001). All panels: n.s., not significant. Mutant comparisons: 5 WT & 5 mutant seedlings/plate. Bar height (mean), error bar (standard error). (**M**) Genetic control of hydropatterning. Auxin establishes a bias in lateral root development between the contact- and air-side of roots. AXR1 modulates bias by regulating the auxin response pathway. Ethylene inhibits LR initiation on the air-side independently of auxin. FLA4 is able to suppress the synthesis of ethylene.

We found that double mutants of *AtFEI1* and *AtFEI2* ( *fei1*/*fei2-1*) as well as single mutants of *AtETO1* (*eto1*) and *AtACS5* (*acs5^eto2^*, which carries a C-terminal mutation in ACS5 that increases the stability of the protein) all showed the same decrease of air-side LR density as observed in the *AtFLA4* mutants *sos5-1* and *sos5-2*. Concomitantly, all mutants showed an increase in contact-side LR density and a reduction in primary root length, as well (Fig. 4, B to D, and fig. S8, C to E). Screening of the single mutants *fei1* and *fei2-1* revealed no significant effects on air-side LR density (fig. S8F), indicating that both genes may act in a redundant fashion, as previously suggested (*35*). The phenotypic similarity of *sos5* alleles with *eto1* and *acs5^eto2^* provides evidence that an increase in ACC production in *sos5* causes the repression in air-side LR development. Indeed, a reduction of ACC synthesis in the *AtACS* hextuple mutant (*acs2-1, acs4-1, acs5-2, acs6-1, acs7-1, acs9-1*) led to a significant increase in air-side LR density (Fig. 4E). We also tested the single mutant *acs5-1*, but observed no difference compared to WT (fig. S8G). This result is likely due to the high-level of redundancy between *ACS* genes in Arabidopsis, which has eight functional ACS homologs (*37*).

Next, we asked whether ACC itself or ethylene, which is synthesized from ACC by ACC-oxidases (ACOs), causes the observed repression in air-side LR development. We found that treatment with 2-aminoisobutyric acid (AIB), a competitive inhibitor of ACOs, caused an increase in air-side LR density and, hence, led to a rescue of the *sos5-2* mutant phenotype (Fig. 4, F and G). Likewise, treatment of Col-0 WT with ACC alone led to a reduction in air-side LR density, likely due to the increased production of ethylene since this effect was reversed by co-treatment with AIB (Fig. 4H). Both ACC and ACC + AIB also showed significant increases in contact-side LR density compared to mock treated Col-0 WT. This increase could be due to a concomitant reduction in primary root length with both treatments (fig. S8H), and/or potential LR inductive effects of ACC itself (*38*). Furthermore, AIB treatment of a strong hydropatterning maize inbred line (33–16) also led to an increase in air-side LR density and a significant reduction of ethylene production (Fig. 4I). Taken together, these observations suggest a central role of ethylene in the suppression of air-side LR development, which does not require localized ACC synthesis, since exogenous ACC application on the contact-side was able to induce the same effect.

We next directly tested the role of ethylene synthesis and perception on hydropatterning. Two *AtACO* quintuple mutants (*ET-free-1* and *ET-free-2*), in which all five *AtACO* genes were mutated by CRISPR/Cas9 (*39*), showed a significant increase in air-side LR density (Fig. 4J). The ethylene perception mutants *ein2-5* and *ein2-50* showed a similar significant increase in air-side LR density (Fig. 4K). Together these observations confirm that ethylene suppresses air-side LR development. Concomitantly, we observed a decrease in contact-side LR density for *ET-free-1*, *ET-free-2*, *ein2-5*, and *ein2-50* that mirrored the increase in air-side LR density leading to no change in total LR density (fig. S8, I and J). Since LR induction can only occur at one of the two xylem poles in Arabidopsis (*40*), it is possible that a derepression of air-side LR development in these mutants leads to a redistribution of LRs from the contact-side. While mutants for *EIN2* have not been described in maize, a mutant allele of *OsEIN2* in rice was available and showed a similar defect in hydropatterning as the Arabidopsis mutant alleles (Fig. 4L), providing further evidence that ethylene-dependent regulation of hydropatterning is conserved between Arabidopsis and grasses.

To investigate whether the ethylene-mediated suppression of air-side LRs is dependent on auxin signaling, we generated a double mutant of *sos5-2* and *axr1-3* since it is known that ethylene induces local auxin biosynthesis (*41*). While the disruption of auxin signaling leads to an increase in air-side LRs, the double mutant *sos5-2*/*axr1-3* showed a significant decrease in air-side LR density (fig. S8K). This observation suggests that each hormone pathway promotes hydropatterning independently.

## Conclusions

Our results reveal that hydropatterning is a crop- and field-relevant response of roots to heterogeneity in soil moisture. Development of modern breeding germplasm in maize led to the weakening of hydropatterning in temperate regions, likely through relaxation of selection. This divergence in hydropatterning may relate to the different selection pressures experienced by each subpopulation. Through genetic analyses, we detected associations between hydropatterning and the auxin and ethylene signaling pathways. Dissection of these pathways in maize, rice and Arabidopsis demonstrates that auxin signaling promotes a bias in LR development towards moisture-contacting surfaces of the root, while ethylene suppresses branching on air-exposed surfaces (Fig. 4M). FLA4, acting at the top of a signaling pathway that restricts ethylene biosynthesis, may perceive an as-of-yet unknown environmental cue to tune the degree to which root architecture is responsive to local soil structure and water availability. A deeper understanding of these pathways may allow for the control of moisture-responsive root growth to improve drought resilience in maize.

## Materials and Methods

### Plant materials

*Zea mays* (maize) inbred lines were obtained from the USDA-ARS National Plant Germplasm System (accession group: Maize.Set.Inbred.Diversity.282.Plus.NAM.Parents) (data S1). Selected seed stocks were bulked up by sib-pollination of plants that were grown in a field at Stanford, CA.

*Arabidopsis thaliana* (Arabidopsis) mutants for orthologs of maize candidate genes were obtained from the Arabidopsis Biological Resource Center (ABRC) (data S11). Additional Arabidopsis mutants that were needed to examine the effect of ethylene on hydropatterning were either obtained from the ABRC or from collaborators and certain crosses were made in house (data S13). For bulking up seeds, selection of homozygous plants, and crosses, Arabidopsis seeds were were surface sterilized for 10 minutes using a 25% (v/v) bleach (Clorox, Product No. 4460032263) and 0.1% Tween® 20 solution (Sigma-Aldrich, CAS No. 9005-64-5) and then washed four times with sterile water. Sterile seeds were then planted on solid media plates with 4.3g/L Murashige & Skoog Salts (Caisson Labs, Ref. No. MSP01-50LT), 1% sucrose (Sigma-Aldrich, CAS No. 57-50-1), 0.05% MES Hydrate (Sigma-Aldrich, CAS No. 1266615-59-1), and 0.4% Gelzan (Sigma-Aldrich, CAS No. 71010-52-1), and stratified for 72 h at 4℃ in the dark. Plates were transferred to a growth chamber and grown under long-day conditions (16 h light / 8 h dark, 22℃, 37% relative humidity) for 10 days after which they were either transferred to soil directly or genotyped for homozygous T-DNA insertions before transfer to soil. Seedlings were grown on PRO-MIX HP soil (Hummert International, Item No. 10200400) to maturity in a growth chamber under long day conditions (16 h light/8 h dark, 22℃, 50% relative humidity). Seeds were collected and stored at room temperature until further use in experiments. Mutant alleles and crosses were confirmed via Polymerase Chain Reaction (PCR) and subsequent analysis via gel electrophoresis. Primers were designed using either the SIGnAL tool (http://signal.salk.edu/tdnaprimers.2.html) or Geneious Prime 2023.2.1 (https://www.geneious.com) (data S11). PCR was performed using the Phire Plant Direct PCR Master Mix Kit (Thermo Fisher, Cat. No. F160S). For mutants with point mutations, subsequent Sanger sequencing was carried out to confirm the genotype of the plants.

### Maize hydropatterning assay

#### Design

The phenotyping system was adapted from *GrowScreen-PaGe* (*42*). With the aim to achieve a uniform moisture gradient across the primary root and hydropatterning of lateral root branches (LRs), we designed a custom-built hydropatterning assay which uses acrylic panels for structural support (fig. S1). In contrast to Gellan-Gum based hydropatterning assays in plates that have been previously described (*9*), our germination-paper based assay is less prone to fungal contamination, since no nutrients are added, and roots have more space to grow. The hydropatterning assay consists of three components (fig. S1A): (1) Rhizo-sheets that were made from laser cut 1/8” black acrylic sheets (Calsak Plastics, Item No. 61299) as a structural support (Sheet.svg). Acrylic sheets were covered on both sides with one layer of pre-moistened Ahlstrom Grade 222 Blot Paper (Great Lakes Filters, SKU: 2228) and one layer of 76# Heavy-Weight Seed Germination Paper (Anchor Paper Company, Item No. SD7630) on the outside. The germination paper was folded twice to form a channel to hold the germinating seeds (fig. S1B). Two small triangular cut-outs were introduced 2.25” from each side at the bottom of the channel to allow the roots to pass through. Sheets were inserted into a pouch made from Phifer BetterVue window screen (Phifer, Item No. 3024751) to prevent roots from growing away from the surface. Each rhizo-sheet was designed to hold up to four seedlings. (2) Rhizo-racks were made from laser cut 1/4” black acrylic sheets (Calsak Plastics, Item No. 61308). Each rack was assembled from three pre-cut sheets (Rack.svg) that were connected using four 3/8“-16 PVC Threaded Rods (McMaster-Carr, Item No. 98871A370) and a total of 24 3/8“-16 PVC Hex Nuts (McMaster-Carr, Item No. 94806A031). Each rack was designed to hold six rhizo-sheets (3) Black polyethylene 12 gallon tanks (Plastic-Mart, Item No. R121812A) were used to hold the rhizo-racks with rhizo-sheets and served as a reservoir for water to keep the filter and germination papers moist (fig. S1C). Each tank was covered with a laser cut 1/8” clear acrylic sheet (Calsak Plastics, Item No. 61413) to reduce water loss by evaporation.

#### Phenotyping

Maize seeds were surface sterilized before germination on 76# Heavy-Weight Seed germination paper (Anchor Paper Company, Item No. SD7630) in 120 x 120 x 17 mm square vented polystyrene petri dishes (Greiner Bio-One, Item No. 688102) for 2 d in a black polyethylene box (Plastic-Mart, Item No. R121812A) in the dark in a walk-in growth chamber (*43*) (fig. S1D). Conditions in the growth chamber were set to 30 °C day / 22 °C night (16-hour photoperiod, 250 μmol m^-2^ s^-1^ light intensity, 50 % relative humidity). In total, we were able to successfully germinate seeds of 285 inbred lines from a total set of 293 inbred lines (data. S1). Once primary roots started to emerge (5 to 10 mm in length), seedlings were transferred into the hydropatterning assay described above and grown for 10 days in the same growth chamber. Positions were randomly assigned with each inbred line being present at least once in each batch. After 10 days, when LRs along the entire length of the primary root on the rhizo-sheets had emerged, 12 cm segments of primary roots were harvested and digitized (contact- and air-side) using a Epson Perfection V800 Photo Scanner (Seiko Epson Corporation, Model B11B223201) with the following settings: Photo Mode, Reflective, 24-bit Color, 1200 dpi, Image Format: TIFF.

#### Image analysis

Images were prepared for counting of LR emergence points using a series of custom macros in ImageJ/Fiji (*44*). First digitized roots were separated into individual files (00_ImageSeparation.ijm) and corresponding air- and contact-side root segments were aligned (01_RegisteredStacks.ijm, 01_Register.py). Then, root segments were traced, measured, and straightened (02_SegmentTrace.ijm). LR emergence points along the contact- and air-side of primary root segments were counted manually using the *Multi-point* tool in ImageJ/Fiji. Counts and coordinates of LRs were exported using the *Analyze* > *Measure* commands.

#### Data analysis and visualization

LR counts from images were imported into R version 4.2.1 (*45*). Data analysis and visualization were performed with the help of the *tidyverse* R package (*46*). Contact- and air-side LR densities were calculated as the sum of all contact- or air-side LRs divided by the length of the root segment that they were counted on (MaizeHydropatterning_PhenoAnalysis.Rmd). Percent air-side LRs was calculated as the sum of all air-side LRs divided by the total sum of all LRs and multiplied by 100. Median and standard deviation were calculated for biological replicate measurements and used for visualization (data S2). After removing roots of inbred lines that showed abnormal growth or had less than 20 LRs on their root segments, we retained phenotypes for 250 inbred lines.

Best Linear Unbiased Prediction (BLUP) values for contact- and air-side LR density as well as percent air-side LRs were calculated by fitting linear mixed models using the lme4 R package (*47*) and treating the box of seedlings as a fixed effect and genotype as a random effect (data S14).

### Microscopy of LR primordia

Maize seedlings were grown in the hydropatterning assay for 3 - 4 days before collecting 7 cm sections of primary roots for microscopy. To be able to distinguish the contact- and air-side after collection, 1 cm^2^ pieces of window mesh were glued with PELCO® Pro CA44 Tissue Adhesive (TED PELLA INC., Prod. No. 10033) to the air-side of the root sections at their distal end.

#### LR primordia quantification

Root segments were fixed in 4% paraformaldehyde (Electron Microscopy Sciences, SKU: 15710) for an hour under vacuum, rinsed four times in 1X phosphate buffered saline (PBS) (VWR International, Item No. MRGF-6235), and transferred to ClearSee [10% Xylitol (Sigma, CAS-No: 87-99-0), 15% Sodium deoxycholate (Sigma, CAS-no: 302-95-4), 25% Urea (Sigma, CAS-No: 57-13-6)] (*48*) with gentle agitation for two weeks. Cleared samples were stained with Basic Fuchsin (Sigma-Aldrich, Item No. S9430) and SYBR GREEN (Sigma-Aldrich, Item No. B0904) (1:1000 1X PBS). Samples were mounted on slides with proper orientation considering the glued mesh piece and imaged on a stereo microscope Leica M205 FCA equipped with 1X PlanApo objective and a fluorescence system with ETGFP and ETDSR filter sets (Leica Microsystems THUNDER Imager Model Organism). Individual frames were stitched together using the Leica LasX Navigator software.

#### Longitudinal sections

To obtain longitudinal sections for detailed images of LR primordia (Fig. 1A), primary root sections were glued to a Vibratome stage using PELCO® Pro CA44 Tissue Adhesive (TED PELLA INC., Prod. No. 10033) and embedded in 8% UltraPure™ LMP Agarose (Invitrogen, Cat. No. 16520-100). 100 μm thick, longitudinal root tissue sections were cut from root tips using a Vibratome Series 1500 (The Vibratome Company, discontinued). Immediately after, sections were transferred to ClearSee [10% Xylitol (Sigma, CAS-No: 87-99-0), 15% Sodium deoxycholate (Sigma, CAS-no: 302-95-4), 25% Urea (Sigma, CAS-No: 57-13-6)] (*48*) for 24 h, then stained with Calcofluor White (Sigma-Aldrich, Item No. 18909) for 60 min and SYBR GREEN (Sigma-Aldrich, Item No. B0904) for 30 min. Finally, sections were mounted on slides and imaged using a Leica SP8 confocal microscope equipped with a Glycerin-immersion 20X objective. Calcofluor White was excited at 405 nm and detected at 425-475 nm while SYBR GREEN was excited at 488 nm and detected at 510-550 nm.

### Correlation analysis

Partial correlation analysis was performed to correlate BLUPs (data S14) of hydropatterning traits (contact-side LR density, air-side LR density, and percent air-side LRs) with BLUPs of phenotypic traits from field-grown maize plants collected in a meta-analysis (*11*). In R, partial Pearson’s correlation coefficients were calculated (MaizeHydropatterning_PartialCor.Rmd) using the *ppcor* library (*49*). Principal components PC1 and PC2 of the population structure (see below) were included as covariates. FDR-adjusted *p*-values were calculated and the significance threshold was set to *p* ≤ 0.05.

### X-ray Computed Tomography imaging

Non-destructive assessment of LR formation in soil was made using a modified version of the method of (*5*). A Newport series loamy sand (sand 83.2%, silt 4.7%, and clay 12.1%; pH 6.35; organic matter 2.93%; FAO Brown Soil) collected from the University of Nottingham farm at Bunny, Nottinghamshire, UK (52.8586°, −1.1280°), air dried and sieved to <2 mm, was packed into PVC columns (52 mm diameter x 150 mm length) to a typical field bulk density of 1.3 g^-1^ cm^3^. Ten replicate columns were saturated with water from the bottom and then allowed to freely drain for two days to notional field capacity. To compare LR formation between soil cores with or without a macropore (to give a scenario where the root has only partial contact with a moist soil surface), a macropore was created from the center of each core using a 7 mm diameter cork borer for half of the replicates. A pre-germinated maize kernel was placed on a 15 mm disk of nylon mesh at the either the center (no macropore samples) or above the macropore and then covered with moist soil taking care not to fill the macropore with soil. The nylon mesh stopped loose soil falling down the macro pore. Plants were subsequently grown for 5 days at 25°C day / 21 °C night (16-hour photoperiod, 250 μmol m^-2^ s^-1^ light intensity, 50 % relative humidity) in a growth cabinet (Conviron, A1000) before X-ray CT scanning. The water content of the soil at field capacity is moist (approximately 26 %), therefore, it is expected that the air within the macropore would have a humid microclimate. A v|tome|x m 240 kV (Waygate Technologies, Germany) X-ray CT system was used to scan each soil core. Scan settings were 160kV X-ray potential energy, 200 µA current, 1 mm aluminum filter on the X-ray tube, 200 ms detector timing, 3000 projection images in FAST scan mode (continuous rotation, no image averaging). Scan resolution was 60 microns with each scan taking 10 mins to acquire. Projection images were reconstructed to 3D volumes using DatosREC software (Waygate Technologies, Germany). LR positions were quantified by manual assessment of the 3D volumes in VGStudioMAX software (Volume Graphics GmbH, Germany). Six strong and weak hydropatterning maize inbred lines were screened (strong: B73, CI64, 33-16; weak: T8, MS153, OH7B).

### Population structure analysis and *Q*_ST_ - *F*_ST_ comparisons

#### Genomic variant selection

Variants for all phenotyped maize inbred lines and selected outgroups were extracted from the *Zea mays* HapMap v3.2.1 data available on the CyVerse Data Commons (*50*). In total 231 maize inbred lines were extracted from the “282_onHmp321” and “hmp321_unimputed” VCF files, respectively, with the vcftools (v0.1.16) ‘--keep’ option (*51*), and merged with bcftools (v1.15.1) ‘merge’ (*52*). We then filtered sites with vcftools to retain only biallelic Single Nucleotide Polymorphisms (SNPs) (--remove-indels --min-alleles 2 --max-alleles 2), remove sites with more than 10% missing data and a minor allele frequency below 0.01 (--max-missing 0.9 --maf 0.01), and prune variants within a 12kb distance from each other (--thin 12000). Since we were interested in keeping only neutrally evolving SNPs to infer population structure and the neutral *F_ST_* distribution, we removed any SNPs falling within the pruning distance from coding regions using bedtools (v2.30.0) ‘window’ (*53*). A distance thinning method was chosen, as pruning based on correlations between SNPs can bias *F_ST_* estimates in structured populations without a homogenous LD distribution, and lead to a more liberal *Q_ST_* - *F_ST_* estimation (*54*). The pruning distance was chosen based on estimates of Linkage Disequilibrium (LD) decay in tropical maize, which drops to r^2^ = 0.2 at a distance of ∼ 4kb and r^2^ = 0.1 at a distance of ∼ 11.6kb (*55*). In total, 74,022 neutral non-coding SNPs were retained for further analysis.

#### Population structure analysis

We examined the population structure in the data set using principal components analysis in PLINK v1.9 (*56*). Genetic clustering was assessed with ADMIXTURE v1.3.0 (*57*) and the results were compared to those of previous studies using microsatellite markers (*8*, *58*). We decided on the final number of clusters by considering the lowest cross-validation error, but also considering what is known about maize subpopulations from the literature. Tropical/subtropical (ts), stiff stalk (ss), and non-stiff stalk (nss) subpopulations were defined by a 0.8 cutoff for the majority ancestry components, while popcorn and sweetcorn were defined a priori due to small sample sizes as previously reported (*8*). Inbred lines without major ancestry components (> 0.8), were defined as mixed.

#### Q_ST_ - F_ST_ comparisons

We calculated global and pairwise *F*_ST_ for the same variants used for population structure analysis, excluding those with a Minor Allele Frequency (MAF) below 0.1. We imposed this additional filter as variants with a low MAF are known to decrease estimates of *F*_ST_ and choosing a higher MAF threshold would therefore make our *Q*_ST_-*F*_ST_ tests more conservative. *F*_ST_ was estimated according to the definition of Weir & Cockerham (*59*).

We converted the VCF file with the vcftools ‘--012’ option to create a genotype matrix and substituted ‘-1’ values for ‘9’ to indicate any missing data. This genotype matrix was used to calculate global Fst with the ‘MakeDiploidFSTMat’ function of the OutFLANK package (*60*) in R. Pairwise *F*_ST_ was calculated with the vcftools (v0.1.16) ‘--weir-fst-pop’ option. Both global and pairwise *F*_ST_ values were weighted prior to comparison with *Q*_ST_ values.

Global and pairwise *Q*_ST_ estimates were calculated for the BLUPs (data S14) of each trait (air-side LR density, contact-side LR density, and percent air-side LRs) using the ‘Pst’ function of the R package Pstat (*61*). A 95% confidence interval was constructed around these estimates by performing 1000 bootstrap replicates using the ‘BootPst’ function of the same package. Distances between the *F*_ST_ and *Q*_ST_ estimates were calculated both globally (once for all populations including mixed individuals, and once excluding mixed individuals) and for each population pair. Differences were considered potentially significant when the confidence interval of *Q*_ST_ did not overlap with the weighted *F*_ST_. To assign levels of significance, we calculated which percentage of the *Q*_ST_ and *F*_ST_ distributions overlapped, with increasing levels from <5% to <1% nand <0.1% of overlap, and corrected the values for multiple testing using the Benjamini & Hochberg correction for all pairwise comparisons (*62*).

### Genome and Transcriptome Wide Association Studies

Genome Wide Association Studies (GWAS) were conducted using the FarmCPUpp (*25*), a C++ implantation of the FarmCPU model (*24*).

Best Linear Unbiased Prediction (BLUP) values for air-side LR density were calculated by fitting linear mixed models using the lme4 R package (*47*) by treating the box of seedlings as fixed effect and genotype as random effect. A total of 1.2 millions of high-density SNPs that were available for 227 of the 250 phenotypes maize inbred lines (*63*) were used as markers for conducting GWAS (MaizeHydropatterning_FarmCPUpp.GWAS.R). Three GWAS runs were conducted using MAF cutoffs ≥ one inbred line (0.4%), ≥ five 5 inbred lines (2.2%), or ≥ 10 inbred lines (4.4%) with the minor allele to identify a spectrum of common to rare trait-associated genetic variants. The first three principal components were calculated using the TASSEL 5.0 software (*64*) and were included as covariates to control for population structure. Trait Associated SNPs (TAS) were selected using false discovery rate (FDR) < 0.05 in R version 4.2.1 (*45*). Linkage disequilibrium between TAS was calculated using PLINK v1.9 (*56*). Multiple TAS that were found in close proximity to each other on chromosomes 3 and 10 showed strong associations between alleles (r^2^ ≥ 0.8, fig. S5D), respectively. Hence, these are likely associated with the same causal variants. Data analysis and visualization were performed with the help of the *tidyverse* R package (*46*) (MaizeHydropatterning_gwasAnalysis.Rmd). For Manhattan plots, the top 40,000 SNPs with the smallest *p*-values were subsampled to create images of a manageable size. Likewise, for quantile-quantile plots, data was subsampled. SNP-genes within 20-kb windows centered on the TAS were identified using B73 version 5 reference gene models (https://download.maizegdb.org/Zm-B73-REFERENCE-NAM-5.0/).

In addition to GWAS, eRD-GWAS (*13*), a Bayesian-based version of Transcriptome Wide Association Study (TWAS) was conducted (MaizeHydropatterning_TWASrun41000.inp) using the RNA-Seq data from root tips of matching 207 maize inbred lines (Kremling et al. 2018). Using an arbitrary Model Frequency threshold ≥ 0.05, significant TWAS-genes were selected in R version 4.2.1 (*45*). Data analysis and visualization were performed with the help of the *tidyverse* R package (*46*) (MaizeHydropatterning_twasAnalysis.Rmd).

### Expression Quantitative Trait Loci

Expression Quantitative Trait Loci (eQTL) mapping for the expression of *Zm00001eb211770* was performed using the FarmCPUpp (*25*), a C++ implantation of the FarmCPU model (*24*). A total of 1.2 millions of high-density SNPs (*63*) of 207 maize inbred lines with matching gene expression data were used as markers for conducting eQTL mapping. The first three principal components were calculated using the TASSEL 5.0 software (*64*) and were included as covariates to control for population structure. Expression associated SNPs (e-SNPs) were selected using false discovery rate (FDR) < 0.05.

Expression associated SNPs (e-SNPs) were selected using false discovery rate (FDR) < 0.05 in R version 4.2.1 (*45*). Data analysis and visualization were performed with the help of the *tidyverse* R package (*46*) (MaizeHydropatterning_eqtlAnalysis.Rmd). For Manhattan plots, the top 40,000 SNPs with the smallest *p*-values were subsampled to create images of a manageable size. Likewise, for quantile-quantile plots, data was subsampled. SNP-genes within 20-kb windows centered on the TAS were identified using B73 version 5 reference gene models (https://download.maizegdb.org/Zm-B73-REFERENCE-NAM-5.0/).

### Hybridization Chain Reaction

Hybridization Chain Reaction (HCR) probes were purchased from Molecular Instruments (www.molecularinstruments.com). Seeds of maize inbred line “B73” were surface sterilized and germinated before transfer to the hydropatterning assay as described above. Once roots had grown in the assay for three days, 2 cm sections of primary root tips were harvested for sectioning directly into FAA fixative (4% formaldehyde, 5% glacial acetic acid, 50% ethanol) on ice. Immediately after, root samples were fixed under vacuum for 1 hour on ice. After fixation, root tips were glued to a Vibratome stage using PELCO® Pro CA44 Tissue Adhesive (TED PELLA INC., Prod. No. 10033) and embedded in 8% UltraPure™ LMP Agarose (Invitrogen, Cat. No. 16520-100). 100 μm thick, longitudinal root tissue sections were cut from root tips using a Vibratome Series 1500 (The Vibratome Company, discontinued). Subsequent sample fixation, dehydration, permeabilization, protease digestion, probe hybridization and amplification was performed as described previously (*65*).

After hybridization and washes, roots with were left in ClearSee [10% Xylitol (Sigma, CAS-No: 87-99-0), 15% Sodium deoxycholate (Sigma, CAS-no: 302-95-4), 25% Urea (Sigma, CAS-No: 57-13-6)] (*48*) overnight at 4 degrees Celsius and then stained with 0.22% Calcofluor-white (Fluorescent Brightener 28, Santa Cruz Biotechnology sc-218504) to visualize tissue layers by staining cell walls. Samples were mounted on glass slides and imaged with a SP8 Leica Confocal Microscope (Leica Microsystems), using a 20X oil immersion objective. For Calcofluor-white stain, excitation was done at 405 nm and collection at 425-475 nm. For imaging of Alexa 647-coupled amplifiers, excitation was performed at 640 nm and collected at 650-675 nm. Leica LAS X Navigator software was used to collect and merge tiled images.

### Arabidopsis hydropatterning assay

#### Phenotyping

Hydropatterning plate assays were performed to identify mutants that showed different hydropatterning behavior compared to Col-0 wild-type (fig. S7A). Seeds were surface sterilized for 10 minutes using a 25% (v/v) bleach and 0.1% Tween solution and then washed four times with sterile water. Sterilized seeds were placed on plates with 4.3g/L Murashige & Skoog Salts (Caisson Labs, Ref. No. MSP01-50LT), 1% sucrose (Sigma-Aldrich, CAS No. 57-50-1), 0.05% MES Hydrate (Sigma-Aldrich, CAS No. 1266615-59-1), and 0.7% Gelzan (Sigma-Aldrich, CAS No. 71010-52-1) and stratified for 72 h at 4℃ in the dark. Each plate contained five Col-0 wild-type seeds and five mutant seeds that were sown about 1 cm from the top. After stratification, plates were then wrapped in parafilm (Fisher Scientific, Cat. No. 13-374-10) with a single layer of micropore tape (VWR International, Cat. No. 56222-182) at the top and placed on a rack to sit vertically in the growth cabinet (Percival Scientific, Model: CU-36L4). Plants were grown under long day conditions (16 h light/ 8 h dark, 22℃, 37% relative humidity) for 11 days. After 11 days of growth, lateral roots were quantified on the contact- and air-side using a S9 E StereoZoom Microscope (Leica, PN: 10 450 814). Lateral roots were classified as “contact-side” when they emerged from the primary root in directions growing on or into the media. All other lateral roots were classified as “air-side”. Subsequently, plates with seedlings were imaged using an Epson Perfection V800 Photo Scanner (Seiko Epson Corporation, Model B11B223201).

#### Image analysis

Root length was measured using a custom built macro (*Arabidopsis_RootTrace.ijm*) in ImageJ/Fiji (*44*).

#### Data analysis and visualization

LR counts from the manual quantification via microscopy and root length measurements from images were imported into R version 4.2.1 (*45*). Data analysis and visualization were performed with the help of the *tidyverse* R package (*46*) (ArabidopsisHydropatterning_MutantScreen.Rmd, ArabidopsisHydropatterning_EthylenePathway.Rmd, ArabidopsisHydropatterning_Pharma.Rmd). Each plate was treated as a biological replicate, since individual seedlings make only few air-side LRs each leading to a large uncertainty when estimating the true frequency of air-side LRs for a genotype. For contact- and air-side LR density, contact- and air-side LR counts were summed up for all Col-0 wild-type and all mutant seedlings per plate and divided by the cumulative root length. Root length was measured from the top most to the bottom most LR counted. Likewise, percent air-side LRs was calculated as the sum of all air-side LRs per genotype/plate divided by the total sum of all LRs per genotype/plate and then multiplied by 100.

To compare the effects of mutants relative to Col-0 wild-type paired Student’s t-tests were used treating each plate as a statistical unit. In cases of comparing treatment effects, plates were still treated as a statistical unit, but non-paired Student’s t-tests were used for comparisons and *p*-values were FDR-adjusted when more than two conditions were compared.

### Pharmacological treatment

#### Arabidopsis

Gellan-Gum plates for the Arabidopsis hydropatterning assay (see above) were prepared with the addition of 2-aminoisobutyric acid (AIB) (Fisher Scientific, Cat. AAA1302114) and/or 1-aminocyclopropane-1-carboxylate (ACC) (PhytoTech Lab, Cat. A1180). Stock solutions (1 M AIB, 10 mM ACC) were prepared in water, filter-sterilized, and added to the media just before pouring the plates. The hydropatterning assay was performed as described above.

#### Maize

Two tanks (*n* = 20 seedlings/tank) with the custom maize hydropatterning assay were prepared as described above. Each tank was filled with 6 L water. One tank was treated with a mock solution while the other tank was treated with a 1 M solution of 2-aminoisobutyric acid (AIB) (Fisher Scientific, Cat. AAA1302114) to a final concentration of 10 mM. Seeds of the strong hydropatterning maize inbred line “33-16” were pre-germinated on plates that were treated either with a mock solution or with 10 mM AIB. Plants were grown, harvested and analyzed as described above.

### Ethylene quantification

Maize root tips were collected for ethylene measurements immediately after harvesting maize plants from the hydropatterning assay. For this, 1cm long root tip sections of four plants were pooled and rolled up in a 1 x 3 cm strip of pre-moistened tissue paper and placed into 2 mL Screw Glass Vials (Thermo Scientific, Cat. No. 6PSV9-1P) filled with 200 μL of water. Ethylene was allowed to accumulate for 29 hours before measurements using a gas chromatograph (Shimadzu, Model GC-8A). From each vial, 1 mL of sample was injected into the gas chromatograph. Ethylene production was normalized by the fresh weight of the root samples.

### Rice hydropatterning assay

Rice (*Oryza sativa*) dehusked seeds of wild-type Nipponbare and *osein2* mutant were surface sterilized for 4 minutes in 100% bleach (CaOCl2 concentration 20%) and then washed 5 times in autoclaved, deionized water. Seeds were sown on plates containing 1/2 strength MS and 1% BD Difco Agar (Fisher Scientific) at a pH of 5.8. Three days post germination, seedlings were transferred to fresh plates and their root tips marked with a dot. Plants were grown in a Conviron growth chamber at 28°C, 70% relative humidity and 150 μmol m^-2^ s^-1^ light intensity at 16-8 hour day-night cycle. All plates were placed in a vertical orientation with an inclination of ∼80° to allow the roots to grow along the surface of ½ MS agar. All hydropatterning measurements were taken from the root growth after transfer (starting from the dot) as primary root orientation might get slightly altered during the transfer process.

Four days after transfer, the plates were imaged and scored for lateral root hydropatterning in a manner similar to that used with Arabidopsis.

## Acknowledgments

We gratefully acknowledge the many thoughtful discussions with Virginia Walbot regarding experiments in maize. We thank the USDA-ARS U.S. National Plant Germplasm System for providing the maize seeds from the Goodman-Buckler association panel. We also thank Carolyn Schultz, the University of Adelaide, and Georg Seifert, BOKU Vienna, for discussions regarding the function and role of FLA4. Thank you to Erick Bautista for managing the greenhouse and field facilities at Stanford University, and all members of the Dinneny laboratory for their help with revisions of the draft manuscript.

## Funding

Faculty Scholars grant from the Simons Foundation and Howard Hughes Medical Institute 55108515 (JRD)

Advanced Research Projects Agency-Energy (ARPA-E), U.S. Department of Energy DE-AR 1565-1555 (JRD)

Biological and Environmental Research (BER) Program, U.S. Department of Energy DE-SC0023160 (JRD)

National Science Foundation MCB-2427432 (JK)

UKRI Frontiers Research ERC StG, EP/Y036697/1 (BKP)

Advanced Research Projects Agency-Energy (ARPA-E), U.S. Department of Energy DE-AR 0000826 (PSS)

Biotechnology and Biological Sciences Research Council (BBSRC)

BB/V003534/1 to (CJS, JB, MB)

Biotechnology and Biological Sciences Research Council (BBSRC)

BB/T001437/1, BB/W008874/1 and BB/W015080/1 (MB)

European Research Council grant HYDROSENSING 101118769 (MB)

HORIZON EUROPE Marie Sklodowska-Curie Actions 765000 (MV)

EVOTREE project RCN 287465 (SB)

## Author contributions

Conceptualization: JDS, JRD

Data acquisition: JDS, TC, AD, CJS, JB, HHT-M, WGV

Data analysis: JDS, ZZ, SB, MV

Visualization: JDS, CJS

Resources: RK, JK, BKP, MB, PSS

Funding acquisition: JRD, JK, BKP, PSS, CJS, JB, MB, MV, SB

Project administration: JDS

Supervision: JDS, JRD

Writing – original draft: JDS, JRD

Writing – review & editing: all authors

## Data and materials availability

Data and summary statistics are available in the supplementary materials. Software, raw data and code for image processing, to generate data, statistics, and figures will be made available on Dryad at the time of publication.

