## Supplemental Figures for "Maize genetic diversity identifies moisture-dependent root-branch signaling pathways"

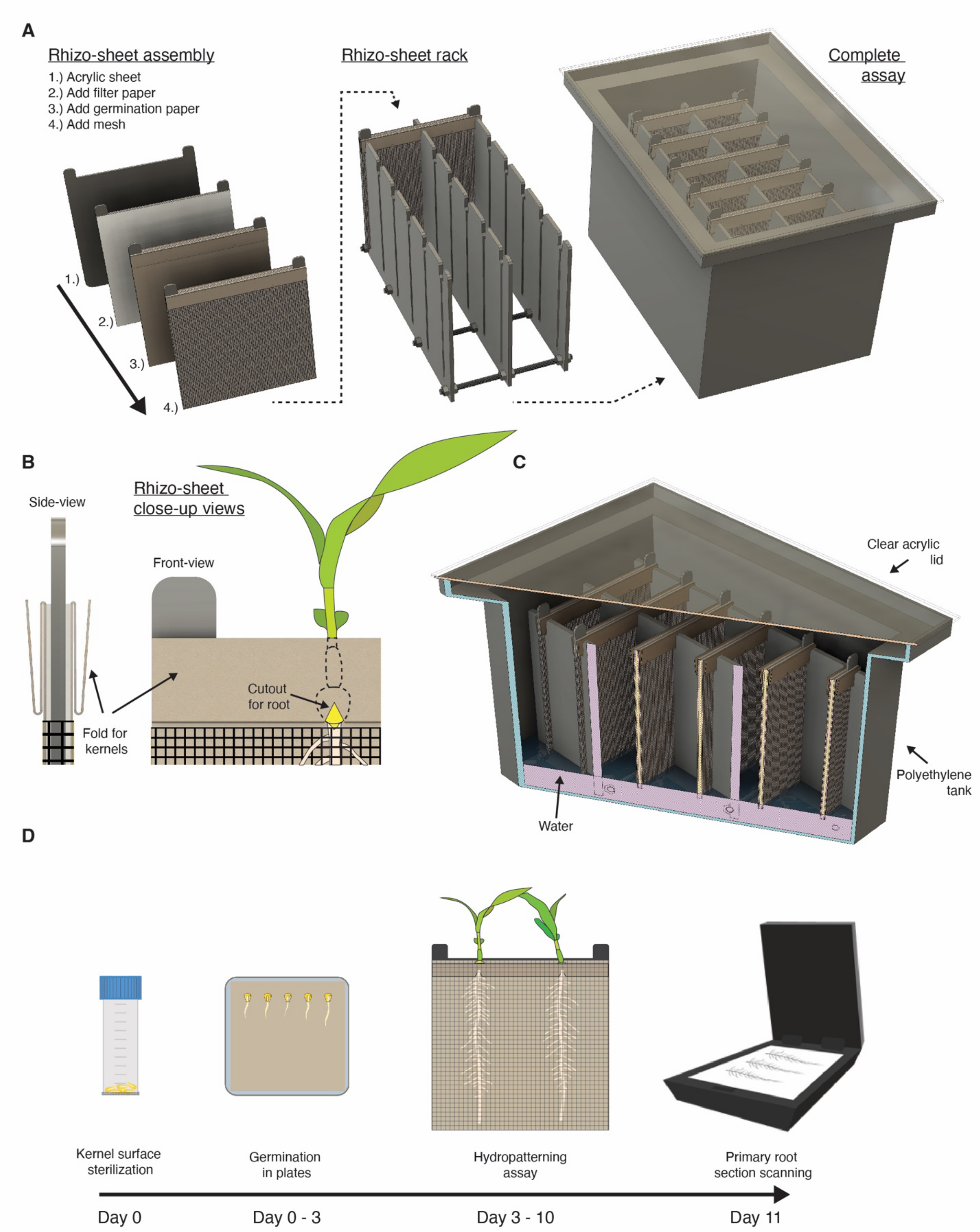


**fig. S1. Design assembly, and workflow of our custom-built assay to characterize hydropatterning in *Zea mays* (maize). (A)** Assembly of the hydropatterning assay: (I) Rhizo-sheets were made from an acrylic panel that was covered on both sides with a layer of filter paper and pre-folded germination paper. The assembly was covered tightly by a layer of window mesh that had been fashioned into a pouch. (II) Rhizo-sheet racks were assembled from laser cut acrylic panels, PVC rods, and PVC nuts and designed to hold six rhizo-sheets each. (III) Polyethylene tanks with clear acrylic lids were used to house the rhizo-sheet racks with rhizo-sheets when running the hydropatterning assay. **(B)** Close-up view of the folded germination paper at the top of the rhizo-sheets. The side-view shows the folded channel to hold germinated maize kernels. The front view shows the triangular cutout that was introduced to let the primary root exit the channel and grow beneath the window mesh along the moist germination paper. **(C)** Cross-section view of the hydropatterning assay showing a 1.5" deep layer of water at the bottom of the tank which kept the rhizo-sheets moist throughout the duration of the experiment. **(D)** Workflow of the hydropatterning assay in maize: (Day 0) Maize kernels were surface sterilized (Day 0 - 3) Sterilized kernels were transferred to plates with moist germination paper and incubated for three days. (Day 3 - 10) Germinated seedlings were immediately transferred to the hydropatterning assay and grown for seven days. (Day 11) 12 cm long primary root sections were cut from the hydropatterning assay and lateral root branch patterns of the contact-and air-side were recorded using a flat-bed scanner.

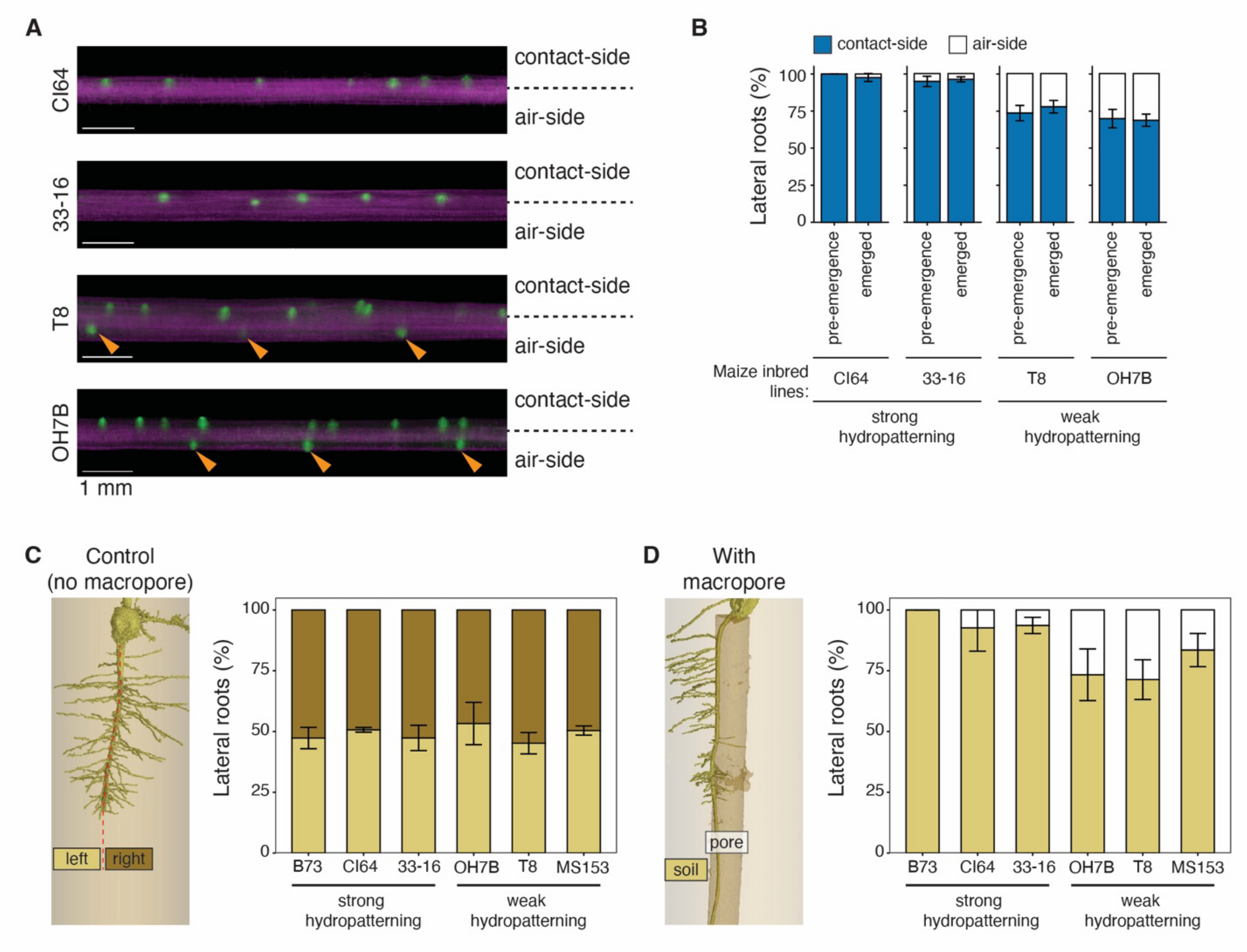


**fig. S2. Microscopy and X-ray Computed Tomography of hydropatterning in *Zea mays* (maize). (A and B)** Fluorescence microscopy and quantification of pre-emergence lateral root (LR) primordia in four maize inbred lines (CI64, 33-16, T8, OH7B). (A) Images show the early mature zone of primary roots for seedlings that were grown in the hydropatterning assay. LR primordia are marked by SYBR GREEN and shown in green. Air-side LR primordia are indicated by orange triangles. White scale bar: 1 mm. (B) Quantitative comparison of pre-emergence LR primordia, assessed via microscopy, and post-emergence outgrown LRs, counted on images from the hydropatterning assay. *n* = 5-10 seedlings/inbred line. **(C and D)** Quantification of LR patterning on primary roots via X-ray Computed Tomography for three strong (B73, CI64, 33-16) and three weak (OH7B, T8, MS153) hydropatterning maize inbred lines. (C) Control experiment with roots fully surrounded by soil. LR emergence points were quantified left and right of the primary root axis. *n* = 2 - 3 seedling replicates/inbred line and n = 36 - 110 LRs/replicate. (D) Primary roots grown through an air-filled macropore in soil. LR emergence points were quantified towards the soil and towards the air-filled macropore. *n* = 2 - 4 seedling replicates/inbred line and n = 13 - 42 LRs/replicate. Bar height (mean), error bar (standard error).

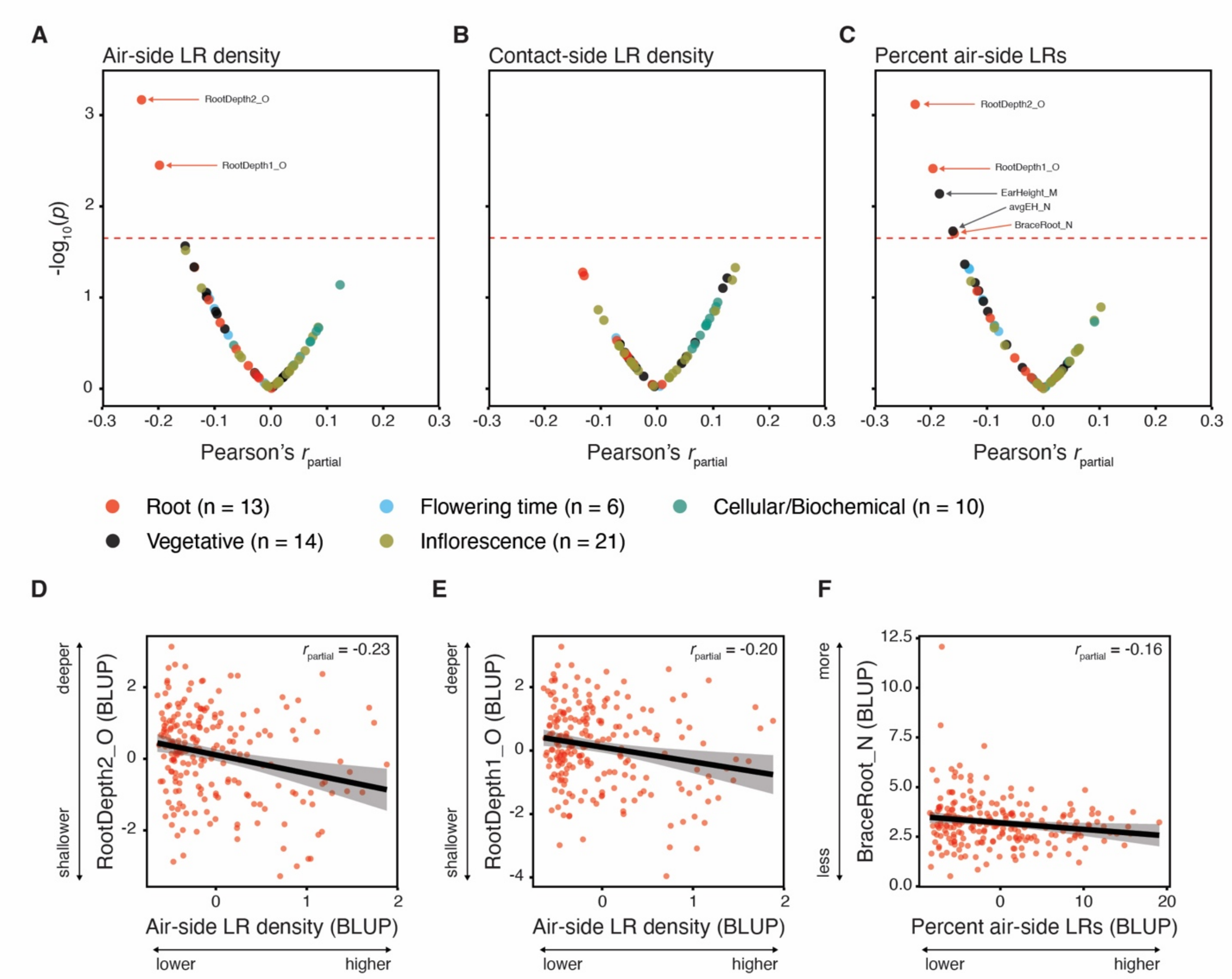


**fig. S3. Correlation analysis of hydropatterning traits with phenotypic traits of field-grown maize plants. (A - C)** Volcano plots for correlations between (A) air-side and (B) contact-side lateral root (LR) density, and (C) percent air-side LRs and 67 quantitative phenotypic trait sets collected from field-grown maize (*11*). Pearson's correlation coefficients were adjusted for population structure ($r_{partial}$). Data points are colored by trait group. Pairwise-complete genotypes (inbred lines) ranged from 195 - 230 per trait set. Red dashed line marks the FDR-corrected threshold at $p = 0.05$. **(D - F)** Partial residuals plot showing the correlations between Best Linear Unbiased Predictions (BLUPs) of (D) air-side LR density and RootDepth2_O, (E) air-side LR density and RootDepth1_O, and (F) percent air-side LRs and BraceRoot_N. RootDepth1_O and RootDepth2_O are measures of root crown depth obtained from excavated root crowns (*12*)*;* $n$ = 218 pairwise-complete genotypes (inbred lines). BraceRoot_N is a measure of the number of nodes with brace roots (*13*); $n$ = 219 pairwise-complete genotypes (inbred lines).Population structure was added as a covariate. Black solid lines (linear regression lines), standard error (gray shaded area).

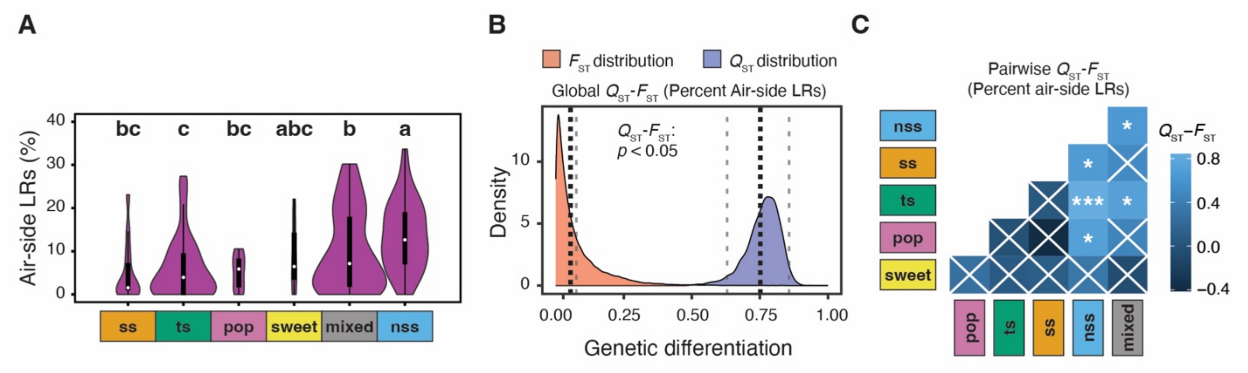


**fig. S4. Phenotypic variation and signatures of selection for percent air-side lateral roots.** (**A**) Comparisons of percent air-side lateral roots (LRs) between subpopulations (ss = stiff stalk, ts = tropical and subtropical, pop = popcorn, sweet = sweet corn, mixed = mixed, nss = non-stiff stalk). Violin plot areas were adjusted by number of inbred lines in each subpopulation ($n_{ss}$ = 14, $n_{ts}$ = 53, $n_{pop}$ = 9, $n_{sweet}$ = 6, $n_{mixed}$ = 88, $n_{nss}$ = 63). Letters denote significant differences between subpopulations ($p \leq 0.05$, Kruskal–Wallis and Dunn's post hoc tests); n.s., no significant differences. (**B**) Population-wide comparison of $F_{ST}$ (fixation index) and $Q_{ST}$ (genetic differentiation in regard to a percent air-side LRs). Black dotted lines denote means, gray dashed lines denote confidence intervals. (**C**) Subpopulation pairwise $Q_{ST}$ - $F_{ST}$ comparisons. Asterisks denote significant differences between $Q_{ST}$ and $F_{ST}$ (* $p \leq 0.05$, ** $p \leq 0.01$, *** $p \leq 0.001$); white crosses, not significant.

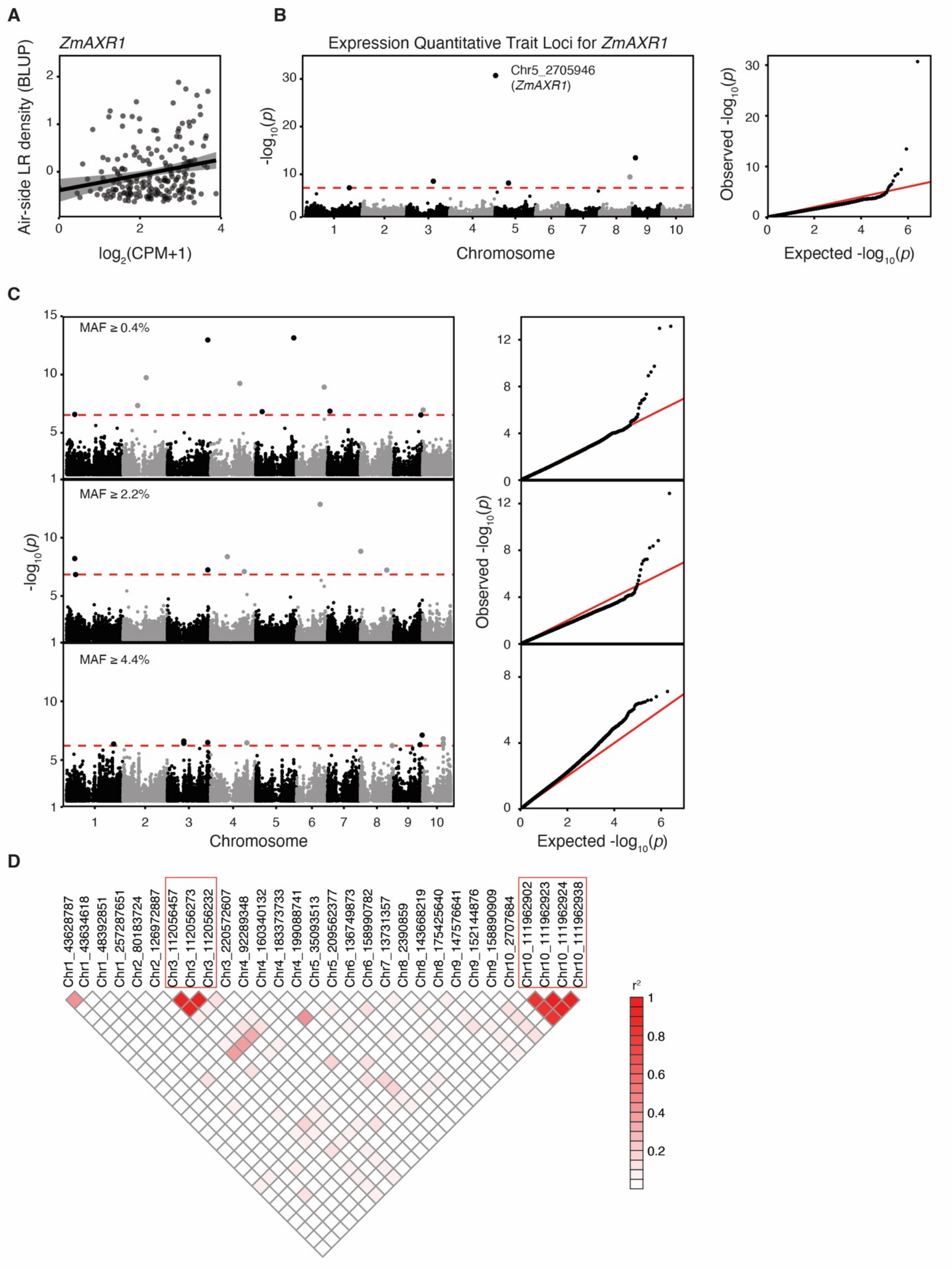
A
ZmAXR1
Air-side LR density (BLUP)
log2(CPM+1)
B
Expression Quantitative Trait Loci for ZmAXR1
Chr5_2705946
(ZmAXR1)
-log10(p)
Chromosome
Observed -log10(p)
Expected -log10(p)
C
MAF ≥ 0.4%
MAF ≥ 2.2%
MAF ≥ 4.4%
-log10(p)
Chromosome
Observed -log10(p)
Expected -log10(p)
D
Chr1_43628787
Chr1_43634618
Chr1_48392851
Chr1_257287651
Chr2_80183724
Chr2_126972887
Chr3_112056457
Chr3_112056273
Chr3_112056232
Chr3_220572607
Chr4_92289348
Chr4_160340132
Chr4_183373733
Chr4_199088741
Chr5_35093513
Chr5_209562377
Chr6_136749873
Chr6_158990782
Chr7_13731357
Chr8_2390859
Chr8_143668219
Chr8_175425640
Chr9_147576641
Chr9_152144876
Chr9_158890909
Chr10_2707684
Chr10_111962902
Chr10_111962923
Chr10_111962924
Chr10_111962938
r2

**fig. S5. Genome and Transcriptome Wide Association Studies for hydropatterning in *Zea mays.*** (**A**) Association between gene expression of *ZmAXR1* (*Zm00001eb211770*) in seedling roots of $n$ = 206 maize inbred lines (*22*) and Best Linear Unbiased Predictors (BLUPs) for air-side lateral rotor density. Black solid line and gray shaded areas show the linear regression line and standard error. (**B**) Manhattan and associated quantile-quantile plot for the mapping of expression Quantitative Trait Loci for *ZmAXR1*. The red horizontal dashed line marks the FDR adjusted significance threshold at $p = 0.05$. (**C**) Manhattan and associated quantile-quantile plots for the Genome Wide Association Study on air-side lateral root density using a total of 1.2 million high-density SNPs (*63*). The three stacked plots show GWAS results for runs at three Minor Allele Frequency (MAF) cutoffs ≥ 0.4%, ≥ 2.2%, ≥ 4.4% which were used to select Trait Associated SNPs (TAS). The red horizontal dashed lines mark the FDR adjusted threshold at $p = 0.05$. (**D**) Linkage Disequilibrium matrix of squared correlations ($r^2$) between all TAS detected by GWAS. Red boxes mark groups of adjacent TAS that are highly ($r^2 \geq 0.8$) correlated.

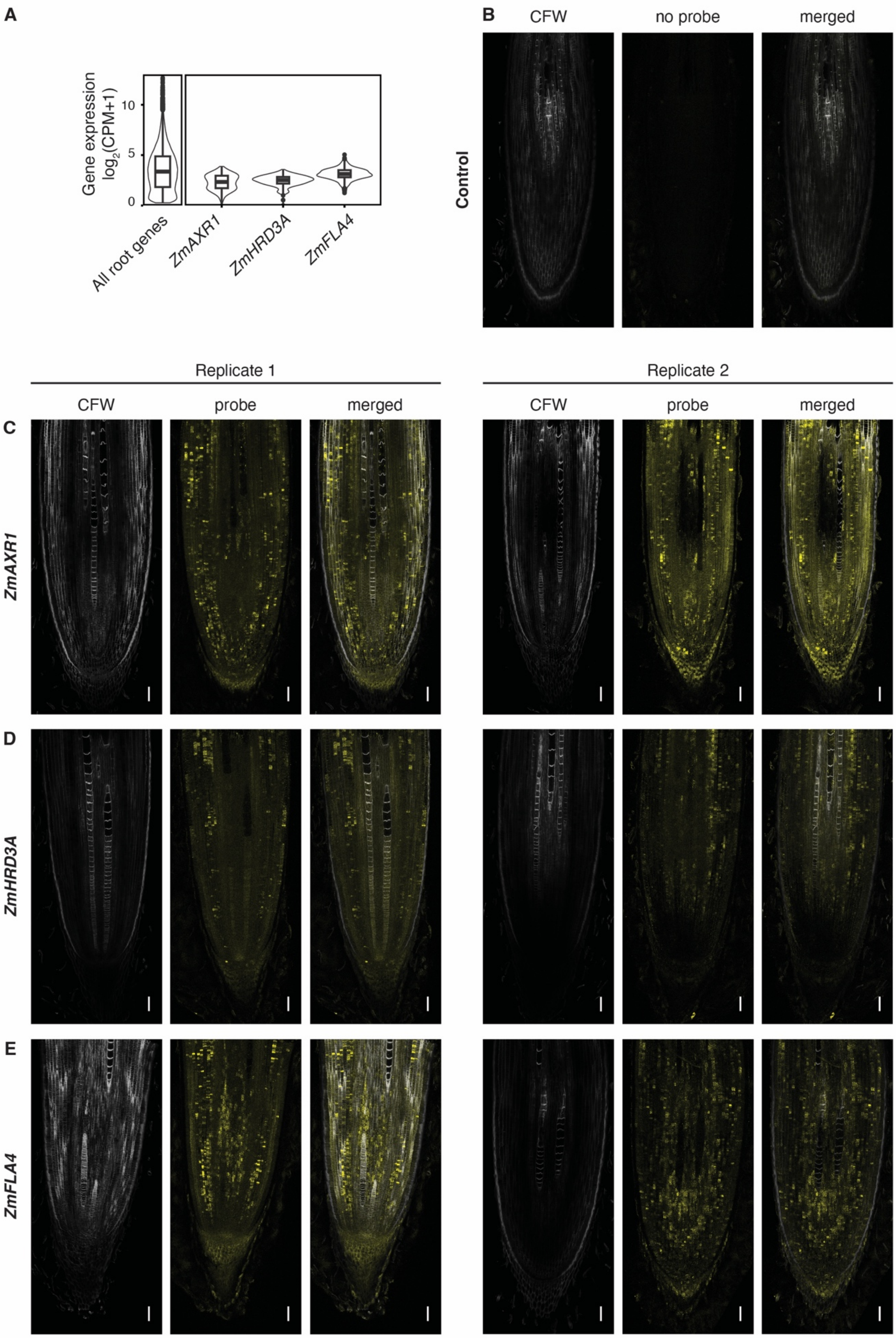

A
B
Gene expression
$\log_2$(CPM+1)
0
5
10
All root genes
ZmAXR1
ZmHRD3A
ZmFLA4
CFW
no probe
merged
Control
Replicate 1
Replicate 2
CFW
probe
merged
CFW
probe
merged
C
ZmAXR1
D
ZmHRD3A
E
ZmFLA4

**fig. S6. Gene expression analysis and Hybridization Chain Reaction (HCR).** (**A**) Gene expression of *ZmAXR1* (*Zm00001eb21177*0), *ZmHRD3A* (*Zm00001eb398230*), and *ZmFLA4* (*Zm00001eb367960*) in root tips of maize seedlings (*22*). Violin and integrated boxplots show the expression across 206 maize inbreds. "All root genes" shows the distribution of the median expression for all root expressed genes (*n* = 21606) across the 206 maize inbred lines. (**B - D**) Image from HCR experiments separated into a channel for Calcofluor White (CFW), a channel for the probe, and a combined channel (merge). (B) Control experiment with no HCR probes added. (C - E) Two independent HCR experiments with probes against (C) *ZmAXR1*, (D) *ZmHRD3A*, and (E) *ZmFLA4*. White scale bars = 100 μM.

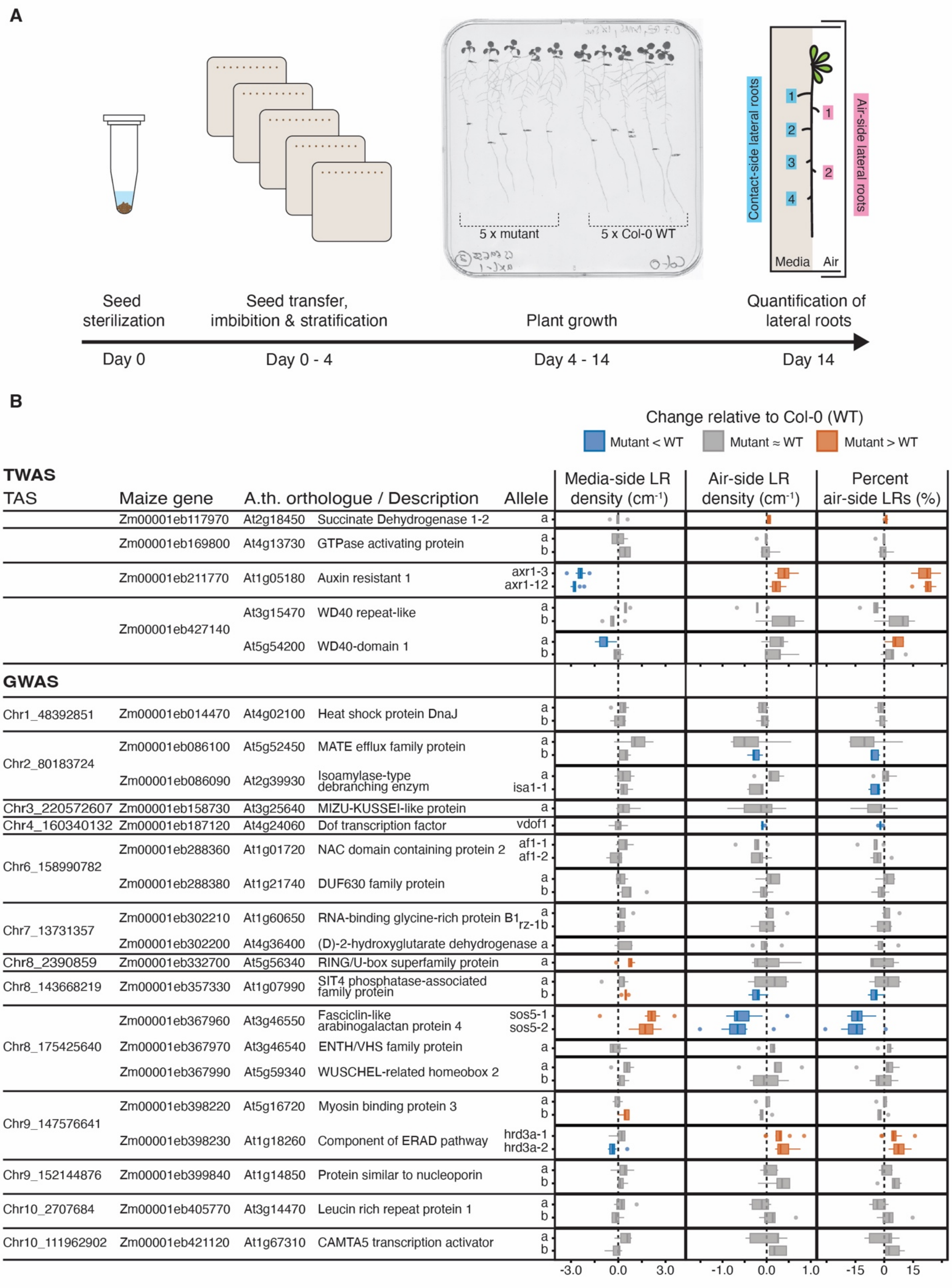

A
5 x mutant
5 x Col-0 WT
Contact-side lateral roots
Air-side lateral roots
Media
Air
Seed sterilization
Seed transfer, imbibition & stratification
Plant growth
Quantification of lateral roots
Day 0
Day 0 - 4
Day 4 - 14
Day 14
B
Change relative to Col-0 (WT)
Mutant < WT
Mutant ≈ WT
Mutant > WT
TWAS
TAS
Maize gene
A.th. orthologue / Description
Allele
Media-side LR density (cm-1)
Air-side LR density (cm-1)
Percent air-side LRs (%)
Zm00001eb117970 At2g18450 Succinate Dehydrogenase 1-2 a
Zm00001eb169800 At4g13730 GTPase activating protein a b
Zm00001eb211770 At1g05180 Auxin resistant 1 axr1-3 axr1-12
Zm00001eb427140 At3g15470 WD40 repeat-like a b
At5g54200 WD40-domain 1 a b
GWAS
Chr1_48392851 Zm00001eb014470 At4g02100 Heat shock protein DnaJ a b
Chr2_80183724 Zm00001eb086100 At5g52450 MATE efflux family protein a b
Zm00001eb086090 At2g39930 Isoamylase-type debranching enzym a isa1-1
Chr3_220572607 Zm00001eb158730 At3g25640 MIZU-KUSSEI-like protein a
Chr4_160340132 Zm00001eb187120 At4g24060 Dof transcription factor vdof1
Chr6_158990782 Zm00001eb288360 At1g01720 NAC domain containing protein 2 af1-1 af1-2
Zm00001eb288380 At1g21740 DUF630 family protein a b
Chr7_13731357 Zm00001eb302210 At1g60650 RNA-binding glycine-rich protein B1 a rz-1b
Zm00001eb302200 At4g36400 (D)-2-hydroxyglutarate dehydrogenase a
Chr8_2390859 Zm00001eb332700 At5g56340 RING/U-box superfamily protein a
Chr8_143668219 Zm00001eb357330 At1g07990 SIT4 phosphatase-associated family protein a b
Chr8_175425640 Zm00001eb367960 At3g46550 Fasciclin-like arabinogalactan protein 4 sos5-1 sos5-2
Zm00001eb367970 At3g46540 ENTH/VHS family protein a
Zm00001eb367990 At5g59340 WUSCHEL-related homeobox 2 a b
Chr9_147576641 Zm00001eb398220 At5g16720 Myosin binding protein 3 a b
Zm00001eb398230 At1g18260 Component of ERAD pathway hrd3a-1 hrd3a-2
Chr9_152144876 Zm00001eb399840 At1g14850 Protein similar to nucleoporin a b
Chr10_2707684 Zm00001eb405770 At3g14470 Leucin rich repeat protein 1 a b
Chr10_111962902 Zm00001eb421120 At1g67310 CAMTA5 transcription activator a b
-3.0 0.0 3.0
-1.0 0.0 1.0
-15 0 15

**fig. S7. Screening of orthologous genes in Arabidopsis for GWAS and TWAS validation. (A)** Workflow of the hydropatterning assay in Arabidopsis: (Day 0) Arabidopsis seeds are surface sterilized (Day 0 - 4) Sterilized seeds are transferred to gel-plates and stratified for three days at 4˚C (Day 4 - 14) Seedlings on plates are grown inside a growth chamber for 11 days (Day 14) Lateral roots (LRs) are quantified using a stereo microscope and plates are imaged using a flatbed scanner for measurements of root length. (**B**) Overview of changes in contact-side LR density, air-side LR density, and percent air-side LRs relative to Col-0 (WT) for 42 Arabidopsis mutant lines. TAS refers to landmark SNP relating to the maize genes identified in through GWAS. $n$ = 5-10 plates/mutant (5 WT & 5 mutant plants/plate). Boxplot fill color indicates statistical differences (Paired Student's t-test, $p \leq 0.05$): significant decrease relative to WT (blue), significant increase relative to WT (orange), and no significant change (gray).

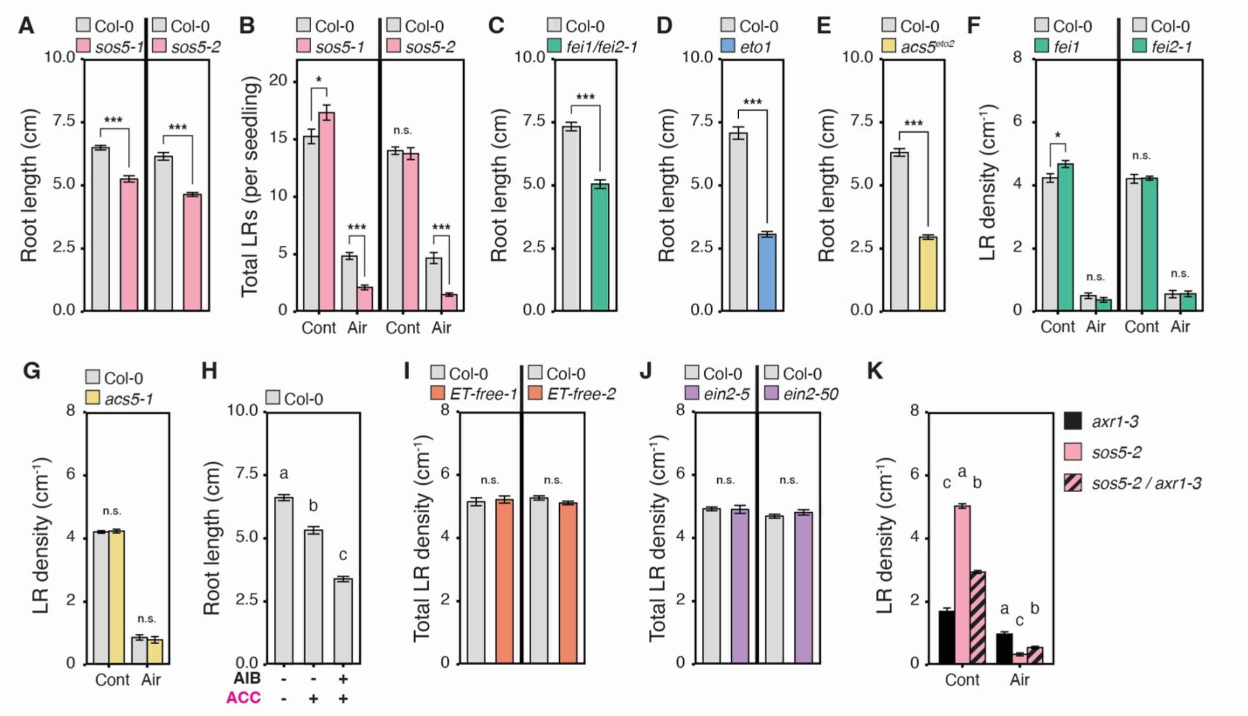


**fig. S8. Additional comparisons of mutants and pharmacological treatments related to the ethylene pathway in Arabidopsis.** (**A, and C - E**) Comparisons of total root length between Col-0 (gray) and (A) *sos5-1* & *sos5-2* (*n* = 10 plates each), (C) *fei1/fei2-1* (*n* = 6 plates), (D) *eto1* (*n* = 5 plates), and (E) *acs5*$^{eto2}$ (*n* = 5 plates). (**B**) Comparisons of the total number of contact-side (Cont) and air-side (Air) lateral roots (LRs) per seedling between Col-0 and *sos5-1* & *sos5-2* (*n* = 10 plates each). (**F and G**) Comparisons of contact-side (Cont) and air-side (Air) LR densities between Col-0 and (F) *fei1* & *fei2-1* (*n* = 6 plates each), and (G) *acs5-1* (*n* = 5 plates). (**H**) Comparisons of total root length between Col-0 treated with a (-) mock solution, (+) 0.05 mM ACC, or (+) 0.05 mM ACC + 5 mM AIB (*n* = 10 plates). (**I, J**) Comparisons of the total number of LRs per seedling (contact- and air-side combined) between Col-0 and (I) *ET-free-1* (*n* = 8 plates) & *ET-free-2* (*n* = 9 plates), and *ein2-5* & *ein2-50* (*n* = 10 plates each). (**K**) Comparisons of Cont and Air LR densities between *axr1-3*, *sos5-2*, and *sos5-2/axr1-3* (*n* = 10 plates each). Asterisks denote significant differences (Paired Student's t-test: * $p \leq 0.05$, ** $p \leq 0.01$, *** $p \leq 0.001$) when Col-0 and mutants were grown on the same plates (5 Col-0 & 5 mutants / plates). Different letters denote significant differences (Student's t-test: $p \leq 0.05$ FDR-adjusted) between treatments or mutant lines when treatments/mutants were on different plates (10 plants/plate). Bar height (mean), error bar (standard error). n.s., not significant.
